# Droplet-based bisulfite sequencing for high-throughput profiling of single-cell DNA methylomes

**DOI:** 10.1101/2023.05.26.542421

**Authors:** Qiang Zhang, Sai Ma, Zhengzhi Liu, Bohan Zhu, Zirui Zhou, Gaoshan Li, J. Javier Meana, Javier González-Maeso, Chang Lu

**Affiliations:** Department of Chemical Engineering, Virginia Tech, Blacksburg, VA 24061, USA; Department of Biomedical Engineering and Mechanics, Virginia Tech, Blacksburg, VA 24061, USA; Department of Pharmacology, University of the Basque Country UPV/EHU, CIBERSAM, Biocruces Health Research Institute, E-48940 Leioa, Bizkaia, Spain; Department of Physiology and Biophysics, Virginia Commonwealth University School of Medicine, Richmond, VA 23298, USA; Department of Genetics and Genomic Sciences, Icahn School of Medicine at Mount Sinai, New York, NY 10029, USA

## Abstract

Genome-wide DNA methylation profile, or DNA methylome, is a critical component of the overall epigenomic landscape that modulates gene activities and cell fate. Single-cell DNA methylomic studies offer unprecedented resolution for detecting and profiling cell subsets based on methylomic features. However, existing single-cell methylomic technologies are all based on use of tubes or well plates and these platforms are not easily scalable for handling a large number of single cells. Here we demonstrate a droplet-based microfluidic technology, Drop-BS, to construct single-cell bisulfite sequencing libraries for DNA methylome profiling. Drop-BS takes advantage of the ultrahigh throughput offered by droplet microfluidics to prepare bisulfite sequencing libraries of up to 10,000 single cells within 2 d. We applied the technology to profile mixed cell lines, mouse and human brain tissues to reveal cell type heterogeneity. Drop-BS will pave the way for single-cell methylomic studies requiring examination of a large cell population.

## Introduction

DNA methylation (i.e., methylation of cytosine at its C5 position) is an important epigenetic mechanism and plays a critical role in regulating genome-wide expression and activity. Diverse DNA methylation profiles are associated with various cell types^1^, developmental stages^2^, and disease states^3^. Single-cell whole genome bisulfite sequencing (scWGBS) is a powerful assay to decipher the heterogeneity of cell-type-specific methylomes at tissue level and genome-wide scale with single-nucleotide resolution. Several tools have been developed to prepare scWGBS library in tubes or well plates, including scBS-seq^4, 5^, snmC-seq^6^, snmC-seq2^7^, sciMET^8^, and sciMETv2^9^. In the early scBS-seq technologies^4, 5^, single cells were isolated into tubes and processed individually without involving barcoding thus only a small number of cells could be studied. snmC-seq^6^ and snmC-seq2^7^ further refined the process toward analyzing a large number of cells by conducting bisulfite conversion in well plates (96 or 384) and using indexed random primers that allowed labeling and pooling of single cells. Finally, sciMET^8, 9^ was developed based on the use of combinatorial indexing to label single cells or nuclei with unique indexes without physical isolation, while the nuclear integrity was maintained. Although a relatively large number of cells (thousands) could be processed using these technologies by running multiple processes in parallel or for an extended period, these platforms do not intrinsically offer high-speed operation.

In contrast, droplet microfluidics allows rapid production and manipulation of millions of water-in-oil droplets as microreactors for isolation and treatment of single cells and has been successfully utilized to produce single-cell sequencing libraries with high throughput for RNA-seq^10, 11^, DNA-seq^12, 13^, ATAC-seq^14^, ChIP-seq^15, 16^, CUT&Tag^17^ as well as multi-omics-seq^18^. In these assays, droplet operations such as encapsulation^10-18^ and fusion^12, 13^ were implemented for physical isolation of single cells or beads and addition of reagents, respectively. Some of the assays^12-14, 18^ also involved molecular biology reaction at elevated temperatures within droplets.

A central mission of single-cell analysis is to reveal heterogeneity and identify subpopulations in tissue samples. The capacity for handling at least 10^4^-10^5^ single cells is crucial for such tasks. For example, a complete analysis of one core biopsy tumor sample requires screening ∼10^4^ single cells^19^. A tissue sample from a specific region of postmortem human brain (e.g. BA9 with direct connections to mental diseases) is typically 0.5 - 1 g in mass, containing roughly 3-6 million neurons and as many glial cells^20^. In this case, survey of 10^4^-10^5^ single cells allows sampling of a small but meaningful fraction of the total cell population.

Here, we demonstrate a high-throughput droplet-based platform for single-cell bisulfite-sequencing (Drop-BS). Our protocol allows production of BS library of 2,000-10,000 single cells within 2 d. In addition to validation using mixed cell lines, we also used the technology to profile various cell types in mouse and human brain samples.

## Results

Drop-BS takes advantage of versatility offered by droplet microfluidics for pairing single cells and barcode beads, mixing various reagents, and conducting multi-step reactions. Briefly, by manipulating single cells and single barcode beads in the droplets, DNA from a single cell was tagged by a unique DNA barcode that was released from a barcode bead (Fig. 1 and Supplementary Fig. 1). The barcoded DNA then underwent bisulfite conversion in droplets. The converted DNA finally underwent random priming and indexing PCR to generate the sequencing library. Drop-BS involves 5 major steps and 3 microfluidic devices to generate Illumina-compatible bisulfite sequencing library from single nuclei (Fig. 1 and 2, Supplementary Fig. 1 and 2). 1) scDNA encapsulation and fragmentation: We used a droplet generation microfluidic device (Fig. 2a) to encapsulate single nuclei and lysis buffer containing micrococcal nuclease (MNase) into droplets (∼35 μm in the diameter) for single-nucleus lysis and genomic DNA fragmentation (Fig. 2c and Supplementary Movie 1). We set the flow rates and concentrations so that ∼10% of these droplets contained single cells and the probability of having double occupancy in a droplet was low (<0.5%). We found that the DNA fragment size affected the final library yield substantially. Generally, short DNA fragments (<150 bp) due to over-digestion were removed by SPRIselect bead selection and under-digestion of DNA yielded less input fragments for barcoding. Thus we optimized the MNase digestion condition (namely CaCl_2_ concentration) to maximize the DNA output (Supplementary Fig. 3). 2) scDNA and single barcode bead droplets pairing and fusion: 3 inlets of the droplet fusion device (Fig. 2b) were used to produce droplets containing single barcode beads and end-repair/ligation reagents (at ∼7.0% bead occupancy; Fig. 2d and Supplementary Movie 2) while 2 side inlets allowed re-injection and spacing of the scDNA droplets generated from step 1 (Fig. 2e and Supplementary Movie 3). We applied alternating current (AC) voltage via an adjacent salt channel to produce dielectrophoresis (DEP)-based fusion^21^ between the scDNA and single-barcode bead droplets (Fig. 2f and Supplementary Movie 4). Averagely ∼80% of the barcode droplets were fused with the scDNA droplets. It is worth noting that there were two depths associated with the channel structures in the droplet fusion device. The depth (as well as the width) of the side channel for re-injection of scDNA droplets was ∼30 μm so that there was no multi-layer stacking of these droplets (Fig. 2e) and simultaneous release of multiple scDNA droplets during fusion. On the other hand, the depth of the rest of the device was ∼70 μm to avoid clogging by barcode bead droplets containing beads of ∼20-40 μm diameter. 3) scDNA ligation and barcoding: The fused droplets were collected into a tube and exposed to UV (Supplementary Fig. 4). The barcoded oligonucleotides were released from barcode beads due to breakage of the photocleavable linker and appended to the fragmented gDNA via ligation reaction in droplets, completing scDNA barcoding. The droplets were broken after barcoding. 4) Droplet-based bisulfite conversion: we discovered that conducting bisulfite conversion in droplets substantially increased the library concentration by a factor of 9 compared to bulk conversion in a tube (Supplementary Fig. 5). We used the bisulfite droplet device (Supplementary Fig. 2) to generate droplets (∼35 μm in the diameter) containing barcoded DNA and bisulfite. The droplets were incubated for bisulfite conversion to achieve a conversion rate of 99.0% (tested using unmethylated lambda DNA as in previous works^7, 8^). The droplets were then broken to pool the DNA. 5) Random priming and indexing PCR: We conducted random priming and indexing-PCR-based amplification to generate the i5/i7-indexed library for sequencing (Supplementary Fig. 6).

**Figure 1.**
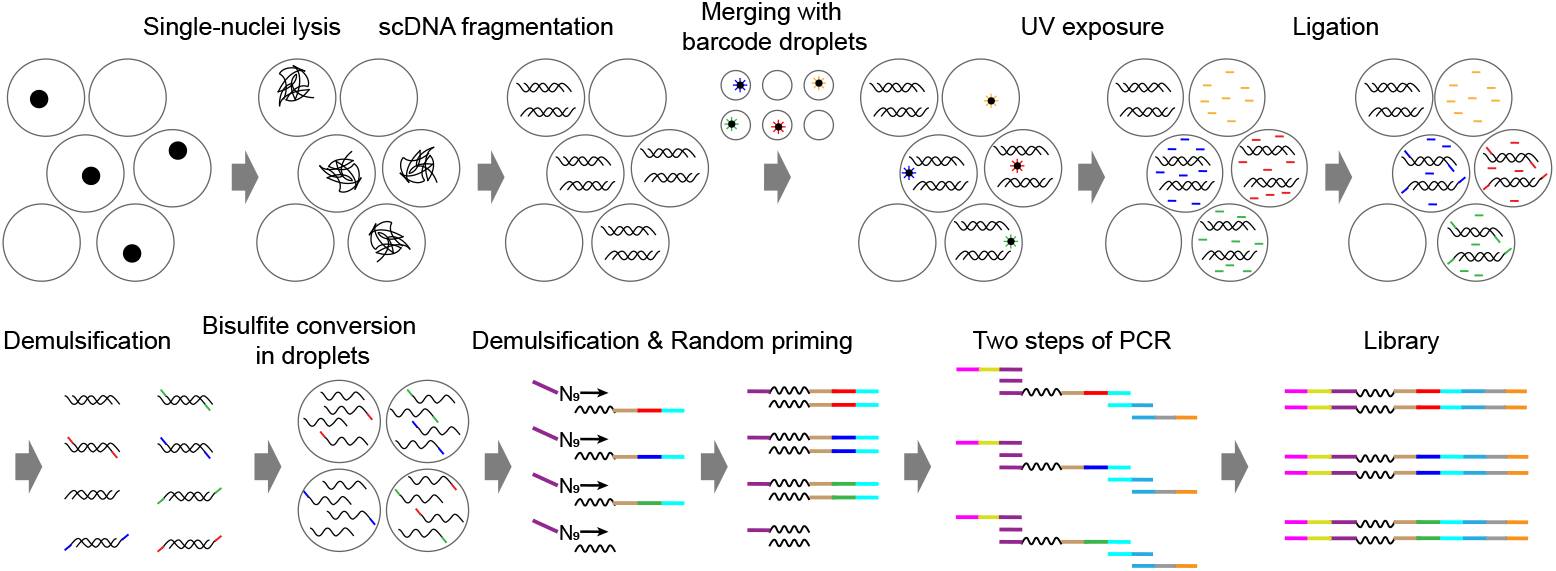
Schematic of Drop-BS library preparation process. Single cells or nuclei are encapsulated into droplets (scDNA droplets) for lysis and DNA fragmentation. After merging scDNA droplets with barcode droplets, fragmented DNA is tagged with barcoded oligos released from barcode beads. Barcoded scDNA is then pooled and further co-encapsulated with bisulfite salt into droplets for bisulfite conversion. Another priming site is added to the DNA using random priming reaction in a tube. Finally, full P5/P7 adapters on the DNA fragments are added by PCR.

**Figure 2.**
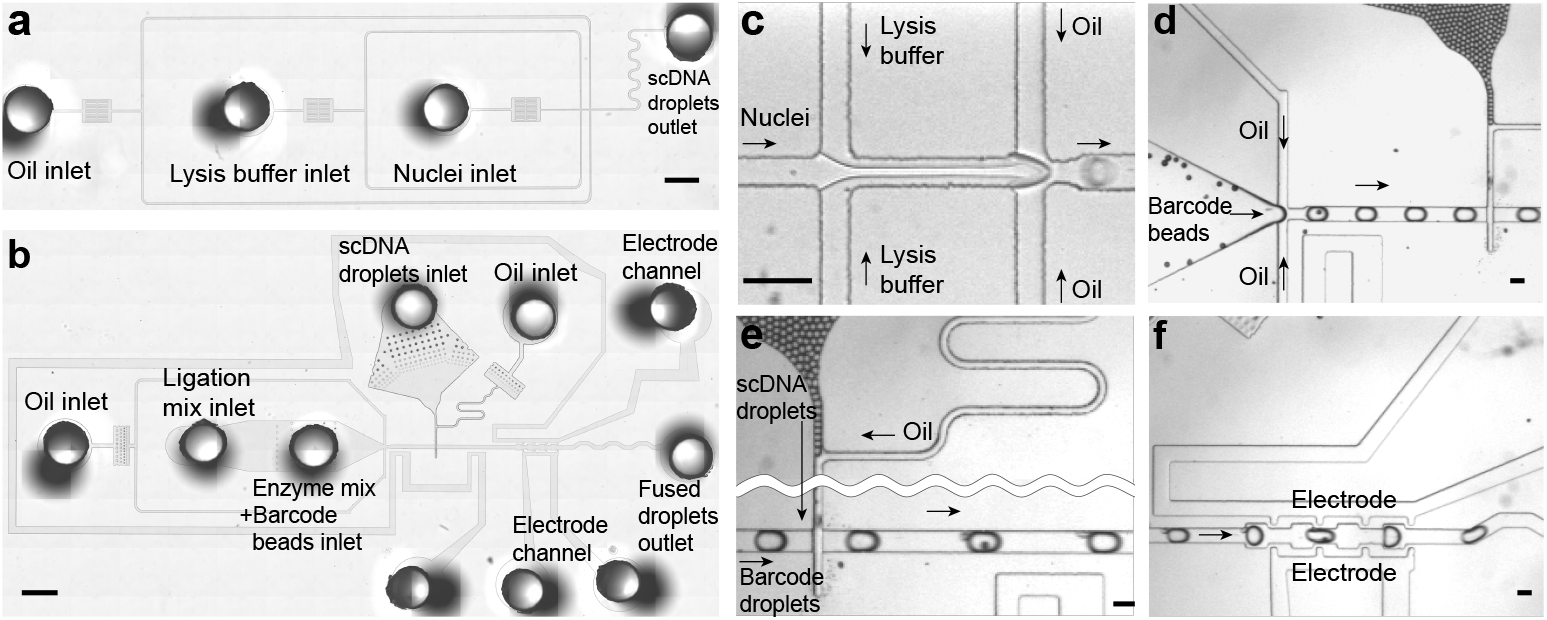
Microfluidic operations of Drop-BS. (a-b) The microscopic images of the droplet generation (a) and fusion (b) devices. The droplet generation device contains three inlets to load oil, lysis buffer, nuclei and one outlet to collect produced scDNA droplets. Barcode bead droplets are produced by a droplet generation module within the droplet fusion device and then paired with reinjected scDNA droplets. There are two channels that are filled with NaCl solution and serve as electrodes for conducting dielectrophoresis-based droplet fusion. One of the electrode channels surrounds most of the microfluidic structure acting as a shield field to prevent reinjected droplets from coalescence. Filtration structures were placed between the inlets and the narrow channels to trap particles and prevent clogging. The device images were created by stitching multiple images. Scale bar: 1 mm. (c-f) Microscopic images of the devices in operation: single-nuclei droplet generation (c); barcode droplet generation in the droplet fusion device (d); scDNA droplet reinjection and pairing with barcode droplets (e); fusion of scDNA droplets and barcode droplets under dielectrophoresis (f). Scale bar: 100 μm.

Drop-BS allows processing a large number of cells or nuclei. Our scDNA droplet generation was conducted with rate of ∼3,000 droplets per second while barcode droplet generation and fusion with scDNA droplets at the rate of ∼132 droplets per second. In our standard operation of processing ∼2,000 cells (involving ∼20,000 barcode beads and ∼574,000 droplets), our operation on the microfluidic devices was finished within a combined 1.3 h, with 10 min on the droplet generation device to produce scDNA droplets, 36 min on the droplet fusion device, and 30 min on the bisulfite droplet device. The entire protocol required 2 d to finish. Up to 10,000 cells can be processed by one operator during the same period (2 d) by increasing the running time on the microfluidic devices or running multiple batches of experiments in the case of the droplet fusion.

We first performed a species-mixing experiment to assess the purity of the single-cell methylomic data generated by Drop-BS. GM12878 nuclei (a human cell line) and mouse brain nuclei were mixed at 1:1 ratio and single-cell bisulfite sequencing library of ∼1,000 cells were prepared. After selection of cell-associated barcodes (Supplementary Fig. 7 and Methods), we recovered 741 high quality barcoded single-cell data, a vast majority (∼96%) of which had >90% of their reads aligned to human hg19 genome (n=445) or mouse mm10 genome (n=266). The data demonstrated that our Drop-BS platform effectively limited crosstalk across droplets and produced high-purity single-cell datasets.

We prepared Drop-BS libraries using three cell lines of different types, as well as a mixture of the three with equal portions. We selected high-quality single-cell data for each sample (GM12878 - 1060 cells, HEK293 - 1489 cells, MCF7 - 1263 cells, and mixed - 1929 cells; See Methods, Supplementary Fig. 8, and Supplementary Table 1). We performed UMAP and Louvain clustering on the methylomic data of the mixture (averagely ∼16,157 unique reads per cell for 1929 cells) and obtained three clusters (Supplementary Fig. 9a). We also conducted co-clustering by adding 300 single-cell data (100 for each cell line) with known identities (Fig. 3a and 3b) to confirm the assignment of the clusters to specific cell lines. We showed that the UMAP clustering was due to different mCG levels associated with the three cell types (Fig. 3c) instead of the number of unique reads per cell (Fig. 3d). However, the unevenness in the number of unique reads among single cells creates spread within the clusters and potentially decreases the resolution of various cell types when they are similar (Fig. 3d). The average global mCG level for each cell type reflected by our Drop-BS data (50.5% for the GM12878 cluster, 68.2% for the HEK293 cluster, and 63.0% for the MCF7 cluster) (Supplementary Fig. 9b and Supplementary Table 1) are similar to previously reported values by bulk assays (48% for GM12878^22^, 66% for HEK293^23^, and 65% for MCF7^24^). Finally, merged single-cell data of the clusters had high correlation with their corresponding merged single-cell data of individually profiled cell lines and previously published methylomic data on the cell lines (Supplementary Fig. 9c).

**Figure 3.**
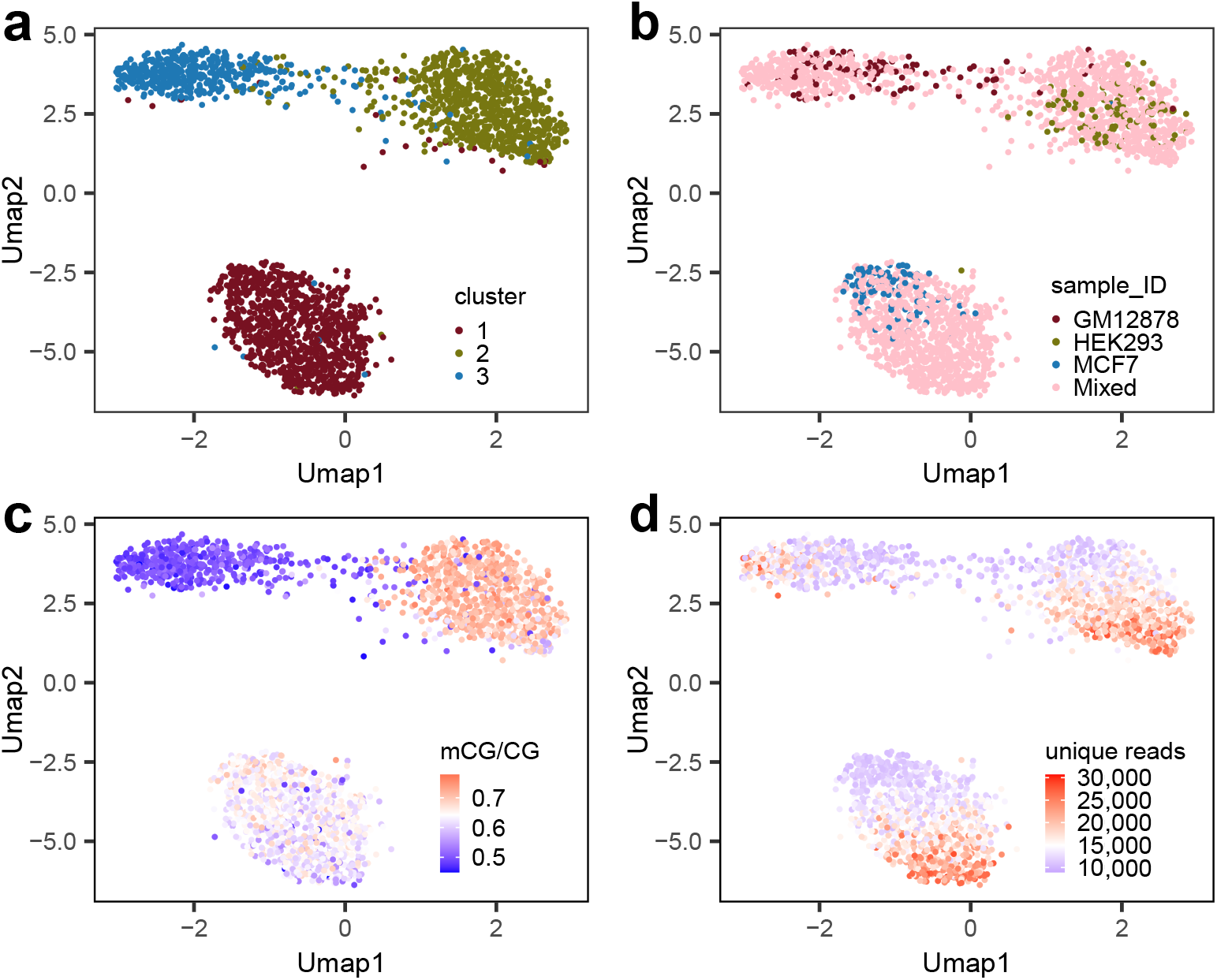
Drop-BS for identifying various cell types in the mixed sample containing three cell lines. (a) Co-clustering of Drop-BS data of the mixed sample (1929 cells) together with 100 single-cell data from each cell line. (b) Co-clustering of the added single-cell data with known identities and the mixed sample yields three clusters. (c) mCG/CG for the single-cell data. (d) The number of unique reads for the single-cell data.

Next, we applied our Drop-BS technology to study prefrontal cortex (PFC) samples from mice and humans. Brain represents the most complex organ and contains a complex mixture of various cell types^6^. The range of CG and CH methylation levels in our Drop-BS data on the mouse and human PFCs were comparable to previously published data (Supplementary Fig. 10). Our Drop-BS data revealed an average global mCG/CG of 71.73% and 76.12%, mCH/CH of 1.85% and 2.71%, for mouse and human PFCs respectively (Supplementary Table 1). The high mCH levels in the brain samples, compared to 1.0% in GM12878 cells, 1.1% in HEK293 cells, and 0.9% in MCF7 cells, are in agreement with literature^25^. We also observed evident hypomethylation regions around TSS and higher methylation level throughout the gene body than that of adjacent intergenic regions (Supplementary Fig. 11).

After discarding low-quality barcodes (See Methods and Supplementary Fig. 8f), we generated 1123 single-cell mouse PFC methylome with averagely 23,932 unique reads per cell and an average mapping efficiency of ∼58% (Supplementary Table 1). Averagely 13,492 CpGs were covered per cell and a total of 11,421,613 CpGs were covered by the merged 1123 single-cell data. We further conducted Louvain clustering of single cell methylomes based on their CH methylation level over genome-wide 100 kb bins. 7 clusters were discovered by Louvain clustering in the mouse PFC sample (Fig. 4a). We then combined the reads from all the cells in each cluster to calculate the CG methylation level over neuron type-specific CG-differentially methylated regions (DMRs) previously profiled by snmC-seq^6^. These neuron type-specific CG-DMRs are regions with lower CG methylation level in a specific cell type than all other cell types^6^. In the mouse PFC sample, we identified that clusters 5 and 7 were excitatory neurons since their CG methylation level in excitatory-neuron-specific CG-DMRs was lower than that in inhibitory-neuron-specific CG-DMRs (Fig. 4b). Clusters 2, 3 and 6 were identified as inhibitory neurons. Clusters 1 and 4 were associated with non-neuronal cells because they showed fairly high methylation rate over most neuron-specific CG-DMRs.

**Figure 4.**
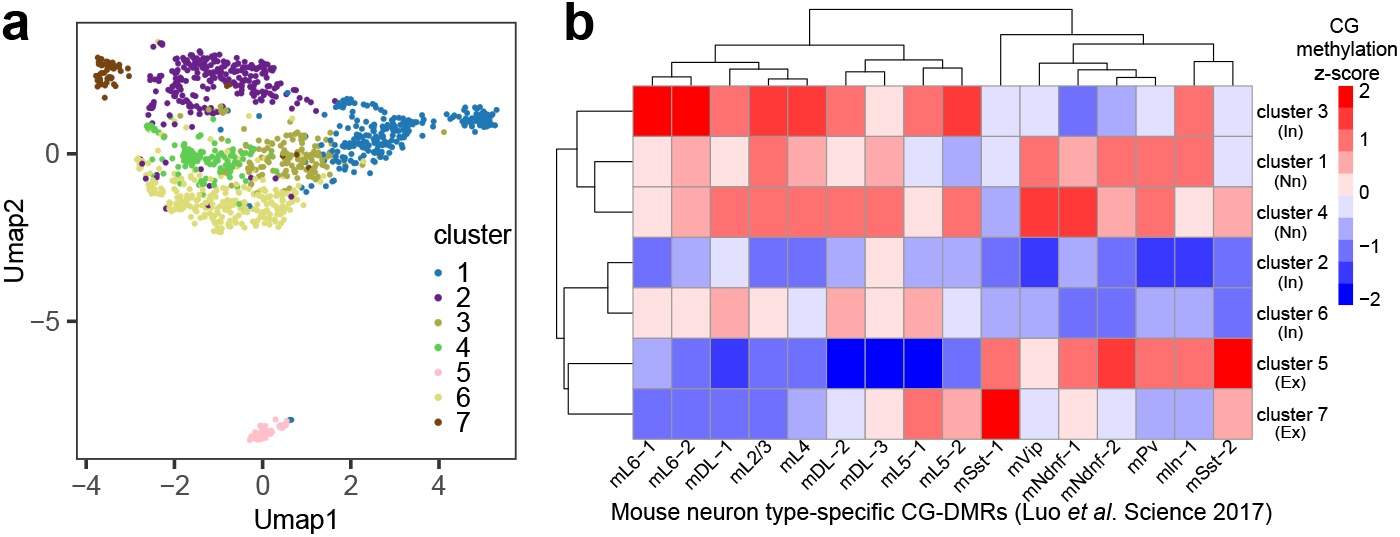
Drop-BS for differentiating cell types based on single-cell CH methylation in mouse brain tissue. (a) Louvain clustering of Drop-BS data on mouse prefrontal cortex. (b) CG methylation z-scores of each mouse PFC cluster over known mouse neuron type-specific CG-DMRs (from Luo *et al*. Science 2017). In: inhibitory neurons; Nn: non-neuronal cells; Ex: excitatory neurons.

We profiled two human brain samples. Human brain 1 (HB1) was from a 41-year-old female containing 1,556 single cells with averagely ∼41,013 unique reads per cell while human brain 2 (HB2) was from a 74-year-old female containing 1,257 single cells with averagely ∼34,630 unique reads per cell (Supplementary Table 1). We first conducted UMAP clustering on the two datasets separately. HB1 Drop-BS data yielded 5 clusters (Fig. 5a) and HB2 data yielded 6 clusters (Fig. 5b). Next, we also combined the two datasets for co-clustering. The two datasets cover similar UMAP space (Fig. 5c). The combined data with a total of 2,813 single cells yielded 7 clusters (Fig. 5d). We confirmed that all the clusters yielded by the individual human brain datasets could mapped to 1-3 clusters among the 7 clusters of the combined dataset (Fig. 5e). 4 out of 5 HB1 clusters and 6 out of 6 HB2 clusters were largely matched to single clusters in the combined dataset (i.e. >53% of single cells in a HB1/2 cluster belonged to a single cluster in the combined data). This demonstrates that the UMAP clustering of our Drop-BS brain data yielded consistent results. Based on CG methylation z-scores over human neuron type-specific CG-DMRs^6^ for the combined clusters, we identified cluster 3 to be inhibitory neurons, clusters 1, 4, 5 to be excitatory neurons, and clusters 2, 6, 7 to be non-neuronal cells (Fig. 5f). We examined the mCH levels at the genome-wide level and in various functional elements for these clusters (Supplementary Table 2). Variation in the mCH level across these clusters were observed in both the global average and on functional elements including genic regions, transcription factor binding sites (TFBS), promoter regions, and CpG islands. Generally, genic regions and TFBS have similar mCH level which was lower than the genome-wide average. Promoter regions presented even lower mCH level than those of genic regions and TFBS, while CpG islands had the lowest mCH/CH among the four types of functional elements. Finally, we conducted pseudobulk differential analysis between the excitatory neuron clusters (1, 4, and 5) and the inhibitory neuron cluster (3) and discovered 637 DMRs (Supplementary Table 3). 594 DMR-associated genes were identified (Supplementary Table 3) and included known glutamatergic or GABAergic neuron markers such as *SATB2, CAMK2A, PROX1, SV2C, GRIK3*, and *SOX6* ^6^. Our Drop-BS data also exhibit expected mCH variation among clusters at these marker genes (Figure 5g).

**Figure 5.**
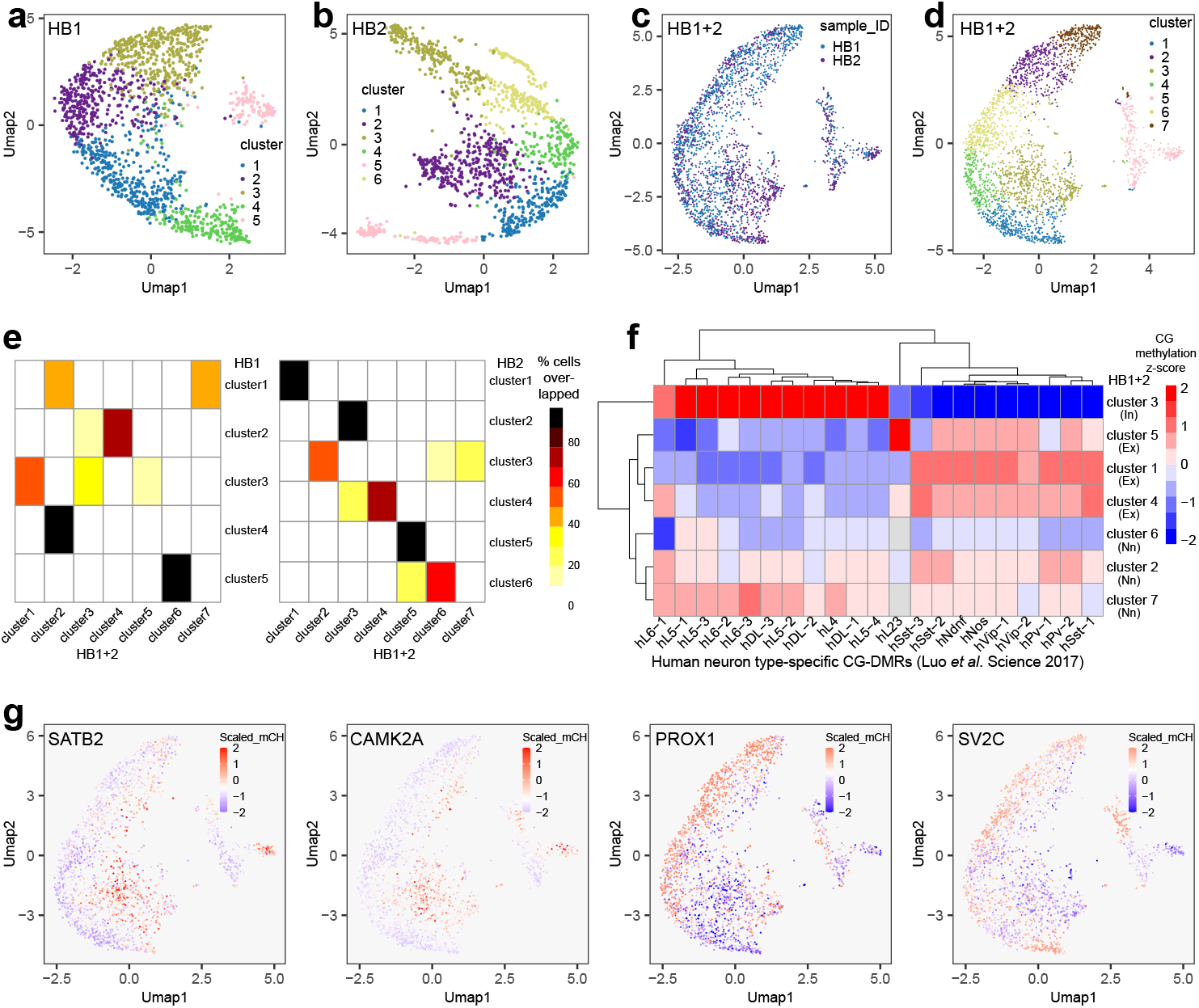
Drop-BS for differentiating cell types based on single-cell CH methylation in human brain tissues. (a-b) Louvain clustering of Drop-BS data on human brain 1 (HB1) and human brain 2 (HB2) prefrontal cortex samples. (c-d) Co-clustering of HB1 and HB2. (e) Comparison of clusters generated on single human brain samples with clusters generated on the combined human brain data. (f) CG methylation z-scores of each HB1+2 cluster over human neuron type-specific CG-DMRs (Luo *et al*. Science 2017). In: inhibitory neurons; Ex: excitatory neurons; Nn: non-neuronal cells. (g) Human marker genes that differentiate inhibitory and excitatory neurons. Single cells are colored according to their normalized mCH level in the gene body.

To summarize, Drop-BS takes advantage of the rapid speed of droplet microfluidics. Droplet generation and fusion occur at a speed of hundreds to thousands of droplets per second and millions of droplets are processed within hours in our protocol. The speed allows producing up to 10,000 single-cell bisulfite sequencing libraries in 2 d. Our droplet-based process offers a bisulfite conversion rate of 99.0%. Drop-BS data achieved an average mapping efficiency of ∼72% for cell lines (n = 5741 cells) and ∼63% for brain samples (n = 3936 cells). The technology provides a high-throughput and scalable approach for studying single-cell methylomes using tissue samples.

## Discussion

Droplet microfluidics has been widely used in high-throughput single-cell genomic profiling^10-18^. Some of these technologies have been successfully commercialized^14, 26^. Here we developed the first high-throughput droplet-based platform, Drop-BS, to construct single-cell bisulfite sequencing libraries for DNA methylome profiling. The challenges for bisulfite sequencing library preparation in droplets lie in genomic DNA fragmentation and bisulfite conversion. DNA fragmentation in bisulfite sequencing was conventionally done by mechanical DNA shearing^27^ which is not compatible with droplet-based assays. In recent single-cell assays, DNA fragmentation was done during bisulfite treatment^6^ and the fragmentation could not be separately optimized independent of the bisulfite conversion process. In Drop-BS, we utilized MNase (Micrococcal nuclease) to randomly cut genomic DNA into fragments within a specific size range by adjusting the treatment condition. We discovered that such optimization was important for increasing DNA recovery. Conventional bisulfite conversion in tubes results in significant DNA degradation^28^. We exploited a strategy of bisulfite conversion in droplets that substantially improved DNA recovery. By conducting pre-bisulfite adapter/barcode tagging, we avoided the complexity of conducting post-bisulfite adapter/barcode tagging^4^ in droplets. Due to the compatibility with other droplet-based single-cell technologies, Drop-BS can potentially be incorporated with other modalities to study multi-omics.

The microfluidic devices used in Drop-BS are compatible with existing commercial platforms (e.g. the ones from Mission Bio). The fabrication follows standard single-layer soft lithography. Barcode beads with designed oligos are readily available from commercial sources. All these features make Drop-BS accessible to a broad community.

In comparison to competing technologies, Drop-BS offers a superb throughput owing to the intrinsic advantage of microfluidic droplet platform but lacks in the number of unique reads yielded per cell (Table 1). With our current protocol, the average number of unique reads per cell plateaus around an assigned sequencing depth of 240,000 reads per cell (Supplementary Fig. 12). Drop-BS data effectively cover important genomic elements such as CpG islands, genic regions, promoter regions, repetitive regions, and TFBS (Supplementary Tables 4 and 5). There is space for further improvement of Drop-BS in terms of its throughput and data quality. First, our droplet generation rate was high (3,000 droplets/sec). Further increase is possible by adjusting the device design and flow rates of aqueous/oil phases. Second, the droplet fusion, required for combination of scDNA droplets and barcode droplets, occurred at a fairly low speed (∼132 droplets/sec) and can potentially be sped up by adjusting the flow parameters while optimizing the DEP conditions to facilitate high-speed fusion. Finally, as it stands, Drop-BS provides a lower number of unique reads per cell compared to competing technologies using well plates or tubes^6, 8^. The underlying reason is possibly the low efficiency of ligation and barcoding reactions within droplets. Systematic optimization of the reaction conditions that are specific to the droplet size and composition will be needed to further improve the data quality.

**Table 1.** Comparison of Drop-BS with other single-cell bisulfite sequencing techniques.

|  | Mapping efficiency | Throughput by one operator | Unique reads per cell (sequencing depth per cell) | CpGs covered per cell | Bisulfite conversion rate |
| --- | --- | --- | --- | --- | --- |
| Drop-BS | 65% | 10,000 cells in 2 d | 41,013 (231,382) | 24,230 | 99% |
| snmC-seq | 55% | 384 cells per wellplate | ~1.7 M (6.9 M) | ~1.7 M | >99% |
| sciMET | 60% | 500-700 cells per experiment | 186,710 (755,529) | 162,045 | 99% |
| sciMETv2 | 69% | ~1000 cells per experiment | 3.1 M (4.9 M) | 2.2 M | 99% |
| scBS-seq | 20% | 44 cells sequenced in the paper | 3.0 M (19.4 M) | 3.7 M | 98.5% |

## Methods

### Cell culture

GM12878 cells were purchased from Coriell Institute and propagated in RPMI 1640 (ATCC, 30-2001) containing 15% fetal bovine serum (Life Technologies Corporation, 16000044), 100 U/mL Penicillin-Streptomycin (contains 100 units/mL of penicillin and 100 μg/mL of streptomycin, Gibco from Thermo Fisher Scientific, 15140122) at 37 °C in a humidified incubator containing 5% CO_2_. GM12878 cells were sub-cultured every two days. HEK293 cells and MCF7 cells purchased from ATCC were cultured in DMEM (high glucose, Life Technologies Corporation, 11965084) containing 15% fetal bovine serum, 100 U/mL Penicillin-Streptomycin at 37 °C in a humidified incubator containing 5% CO_2_. HEK293 cells and MCF7 cells were sub-cultured when reaching ∼80% confluency.

### Mouse brain samples

C57BL/6J mice were purchased from Jackson Laboratory and maintained with 12-h light/12-h dark cycles and food and water ad libitum. 10-week-old male mice were sacrificed by compressed CO_2_ followed by cervical dislocation. Mouse brains were rapidly dissected, frozen on dry ice, and stored at -80°C. To collect prefrontal cortex (PFC) samples, mouse brains were sectioned coronally at Bregma (1.90 to 1.40 mm) with a razor blade in dissection media (20 mM Sucrose, 28 mM D-Glucose, 0.42 mM NaHCO_3_ in HBSS). The dissected brain tissues were stored at -80 °C until use. This study was approved by the Institutional Animal Care and Use Committee (IACUC) at Virginia Tech.

### Human brain samples

Human brains were obtained during autopsies performed at the Basque Institute of Legal Medicine, Bilbao, Spain. The study was developed in compliance with policies of research and ethical review boards for post-mortem brain studies (Basque Institute of Legal Medicine, Spain). Specimens of frontal cortex (Brodmann area 9) were dissected at autopsy (0.5-1 g tissue) on an ice-cooled surface and immediately stored at -80°C until use. Tissue pH values were within a relatively narrow range (6.7 ± 0.08). Brain samples were also assayed for RIN (RNA integrity number) values (7.87 ± 0.21). The brain tissue for the human brain 1 (HB1) sample belonged to a 41-year-old Caucasian female and the brain tissue for the human brain 2 (HB2) sample belonged to a 74-year-old Caucasian female with a postmortem interval (PMI) of 22 h, negative medical information on the presence of neuropsychiatric disorders or substance use disorder, and negative toxicological screening results for either psychotropic drugs or ethanol.

### Nuclei extraction from mouse/human brain tissues

To isolate nuclei, the PFC was placed in 5 ml ice-cold nuclei extraction buffer (0.32 M sucrose (Sigma-Aldrich, S0389-500G), 5 mM CaCl_2_(Sigma-Aldrich, 21097-50G), 3 mM Mg(Ac)_2_ (Sigma-Aldrich, M5661-50G), 0.1 mM EDTA (Life Technologies, 15575-038), 10 mM Tris-HCl (Life Technologies, 15568-025), and 0.1% Triton X-100 (Sigma-Aldrich, T8787-100ml) with freshly added 50 μl of PIC (Protease inhibitor cocktail; Sigma-Aldrich, P8340), 5 μl of 100 mM PMSF (Phenylmethanesulfonyl fluoride; Sigma-Aldrich, P7626-1G) and 5 μl of 1M DTT (Sigma-Aldrich, 43816-10ml)). The tissue was thawed and homogenized by slowly douncing on ice with a loose pestle A (Kimble, 7-ml size) for 15 times, and with a pestle B for 25 times. We filtered the homogenate with a 40-μm nylon mesh cell strainer (Fisher Scientific, 22-363-547) and then centrifuged the sample at 1,000 g for 10 min at 4 °C. We then removed the supernatant and resuspended the pellet in 500 μl ice-cold nuclei extraction buffer.

The suspension was mixed with 750 μl of 50% iodixanol solution (50% iodixanol (Sigma-Aldrich, D1556-250ml), 25 mM KCl (Sigma-Aldrich, P9333), 5 mM MgCl_2_ (Sigma-Aldrich, M2670-100G), and 20 mM Tris-HCl (pH 7.8)) and centrifuged at 5,000 g for 15 min at 4 °C. After centrifugation, we followed the s3_WGS protocol^29^ to perform nuclei fixation and nucleosome depletion. Briefly, the nuclei pellet was resuspended in 5 ml of NIB:HEPES buffer (10 mM HEPES-KOH (pH 7.2; Sigma-Aldrich, H0527), 10 mM NaCl (Sigma-Aldrich, 7647-14-5), 3 mM MgCl_2_, 0.1 % Igepal (Sigma-Aldrich, I8896-50ML), 0.1 % Tween-20 (Sigma-Aldrich, P9416-50ML)) in a 15 ml centrifuge tube (Corning, 430791). We incubated the mixture at room temperature for 10 min on a rotator (speed: 50 rpm, Argos Technologies RotoFlex) after adding 246 μl of 16 % formaldehyde (Thermo Fisher Scientific, 28908) to the suspension. The mixture was centrifuged at 500 g for 5 min at 4 °C to form nuclei pellet and the supernatant was then removed. 1 mL of NIB:Tris (10 mM Tris-HCl (pH 7.4), 10 mM NaCl, 3 mM MgCl_2_, 0.1 % Igepal, 0.1 % Tween-20) was then added to resuspend and wash the nuclei pellet to quench the fixation. The nuclei were pelleted by centrifuging at 500 g for 5 min at 4 °C and resuspended in 200 μl 1X NEBuffer 2.1 (50 mM NaCl, 10 mM Tris-HCl, 10 mM MgCl_2_, 100 μg/ml BSA; NEB, B7202) in a 1.5 ml tube. The suspension was centrifuged at 500 g for 5 min at 4 °C to pellet nuclei which were then resuspended in 760 μl of 1× NEBuffer 2.1. We added 40 μl of 1 % SDS (Life Technologies,153-035) to the nuclei suspension and incubated the mixture at 37 °C for 20 min on a Multi-Therm shaker (Benchmark Scientific, H5000-HC) with a shaking speed of 300 rpm. After SDS treatment, the mixture was centrifuged at 500 g for 5 min at 4 °C. The pellet was resuspended in 1 ml of ATAC-RSB buffer (10 mM Tris-HCl (pH 7.4), 10 mM NaCl, 3 mM MgCl_2_ with freshly added 10 μl of 10% Tween-20). The suspension was pipetted up and down several times and then centrifuged at 500 g for 5 min at 4 °C. Finally, the nuclei were resuspended in 100 μl of ATAC-RSB buffer with freshly added 1 μl of 10% Tween-20. We measured the concentration of the nuclei using a hematocytometer and diluted the nuclei concentration to 5 million/ml using the ATAC-RSB buffer.

### Nuclei extraction from cell lines

1 ml suspended cells were centrifuged at 300 g for 5 min at 4 °C. Cells were then resuspended in 500 μl ice-cold nuclei extraction buffer. The rest of the procedure was identical to the process described above for brain tissues.

### Microfluidic device fabrication

Three microfluidic devices were designed and fabricated in this work using soft lithography^30, 31^, including droplet generation device, droplet fusion device, and bisulfite droplet device. The channel structure was designed using LayoutEditor (juspertor GmbH, Germany) and printed on a transparency photomask with high resolution (10,000 dots per inch, Fineline imaging).

To produce the masters for molding of the droplet generation device (Fig. 2a) and the bisulfite droplet device (Supplementary Fig. 2), one photomask was used for each. A layer (∼30 μm in thickness) of SU-8 2025 photoresist (MicroChem, Y111069 0500L1GL) was coated on a 3-inch silicon wafer (University Wafer, P(100)1-100ohm-cm SSP) by spinning at 500 rpm for 10 s followed by 3000 rpm for 30 s. The wafer was then soft baked at 95 °C for 7 min. The wafer with coated photoresist was covered by the photomask (Supplementary Data Set 1 and Supplementary Data Set 2; the files can be opened by LayoutEditor (https://layouteditor.com/).) and then exposed to 365 nm UV (160 mJ/cm^2^) for 17 sec. The wafer was baked at 95 °C for 7 min and then rinsed with SU-8 developer (MicroChem, Y0201004000L1PE). The wafer was finally washed with isopropanol, dried in nitrogen stream, and baked at 150 °C for 15 min to produce a master for device molding.

To produce the master for molding of the droplet fusion device (Fig. 2b), two photomasks (Supplementary Data Set 3) were used to generate features of different thicknesses on the same wafer. First, a layer of SU-8 2025 photoresist with ∼70 μm thickness was coated on a 3-inch wafer by spinning at 500 rpm for 10 s followed by 1250 rpm for 30 s. The wafer was then soft baked (95 °C for 7 min) and exposed to 365 nm UV (160 mJ/cm^2^) for 17 sec while covered by the first photomask (Supplementary Data Set 3 lower). The wafer was baked again (95 °C for 7 min). After rinsing with SU-8 developer, another SU-8 2025 photoresist layer of ∼30 μm thickness was coated on the same wafer by spinning at 500 rpm for 10 s followed by 3000 rpm for 30 s. The wafer then underwent soft baking (95 °C for 7 min), and UV exposure (160 mJ/cm^2^, 365 nm for 17 sec) under the second photomask (Supplementary Data Set 3 upper). The second photomask was visually aligned with the features created by the first photomask before exposure. The wafer was baked (95 °C for 7 min) after exposure. The wafer was then developed in SU-8 developer, followed by rinsing with isopropanol, drying by nitrogen, and baking at 150 °C for 15 min to generate the master.

The devices were molded in polydimethylsiloxane (PDMS). Degassed PDMS prepolymers (Dow Corning, Sylgard 184) mixed with catalyst at a ratio of 10:1 (w/w) was poured onto a master, degassed again, cured at 80 °C for 1.5 h. The cured PDMS slab with a ∼5 mm thickness was slowly peeled from the master. Inlet and outlet holes were punched into the PDMS slab. The PDMS slab and a pre-cleaned glass slide were then placed in a plasma cleaner (Harrick Plasma, PDC-32G) and treated for 1 min. The oxidized PDMS and glass surfaces were brought into contact and bonded to form a permanent seal after baking at 80 °C for 24 h. The microfluidic devices were rendered hydrophobic prior to use by treating with Aquapel (Aquapel Glass Treatment, PGW Auto Glass, LLC) for 2 min followed by baking at 80 °C for 10 min.

### Design and preparation of barcode beads

Barcode beads with customized oligonucleotide sequences (LOT #: L142464-1) were purchased from ChemGenes (Wilmington, MA). Specifically, each oligonucleotide on the bead surface consisted of four parts (Supplementary Fig. 1): (1) a photo-cleavable linker that connects the oligonucleotide to the bead surface and releases the oligonucleotide after UV exposure; (2) a PCR priming site (identical for all beads); (3) a 15-nt barcode sequence of A, T, G bases for labeling single cells (unique for each bead); (4) a ligation site.

To form a double-stranded tail for ligation, we annealed a complimentary 19 nt-oligonucleotide (called “3’-end blocked complement oligo”, ordered from IDT with the sequence shown in Supplementary Fig. 1) to the bead oligonucleotides. Briefly, ∼20,000 barcode beads were washed three times with 0.5 ml of TE-TW (10 mM Tris pH 8.0, 1 mM EDTA, and 0.01% Tween-20). We mixed 70 μl oligo annealing buffer (10 mM Tris pH 8.0, 50 mM NaCl, and 1 mM EDTA) and 30 μl of 100 μM complimentary oligonucleotide and resuspended the barcode beads in it. We incubated the mixture at 85 °C with a shaking spend of 1500 rpm for 5 min on a Multi-Therm shaker (Benchmark Scientific, H5000-HC). Then we let it slowly cool down to room temperature while continuing to shake the mixture at 1500 rpm. The prepared barcode beads were washed with TE-TW three times prior to loading into the microfluidic device.

### Single-nuclei lysis and DNA fragmentation in droplets

In a typical experiment, we used the below protocol to generate bisulfite sequencing libraries on ∼2,000 single cells. Three 0.5 m-long PFA tubing (IDEX Health & Science, 1622L) connected to three syringes were loaded with 100 μl of the nuclei suspension, 160 μl of 2x lysis buffer (2 % Triton x-100, 100 mM Tris-HCl pH 8.0, 300 mM NaCl, 0.2% sodium deoxycholate (Sigma-Aldrich, 302-95-4), 10 mM sodium butyrate (Sigma-Aldrich, B5887-1G), 0.375 mM CaCl_2_, 0.0375 U/μl MNase (Thermo Fisher Scientific, 88216), 0.2% Sarkosyl (Sigma-Aldrich, L7414), with freshly added 1.6 μl of PIC and 1.6 μl of 100 mM PMSF), and 1 ml oil (Bio-Rad, 1864006), respectively. The three syringes were mounted on three separate syringe pumps (Chemyx, FUSION 200) and the reagents were flowed simultaneously into the three inlets of the droplet generation device (Fig. 2a) at the flow rates of 2 μl/min for the nuclei suspension, 2 μl/min for the lysis buffer, and 14 μl/min for the oil to generate droplets with a size of ∼35 μm in diameter. We continuously collected the generated droplets into a 0.2 ml tube that was placed on ice for 10 min to gather about 1.78 million droplets. We incubated the droplets on ice for 10 min, then at room temperature for 15 min, and finally at 65 °C for 30 min. After incubation, excessive oil at the bottom of the tube was removed and the tube was put on ice until use.

### Single-cell DNA barcoding in droplets

The electrode channels (Fig. 2b) in the droplet fusion device were filled with 2 M NaCl solution and connected to a high voltage alternating current (AC) power supply with an output set at 300 Vp-p and 200 kHz (Micromechatronics, PDUS210-800V). Single-cell droplets generated after cell lysis and DNA fragmentation were re-injected into the droplet fusion chip (Fig. 2b) at a flow rate of 0.12 μl/min. About 20,000 prepared barcode beads were resuspended in 40 μl enzyme buffer (1 U/μl Fast-Link DNA Ligase (Lucigen, E0077-2-D3), 20% End-It Enzyme Mix (Lucigen, E0025-D1), 18% Glycerol) and flowed into the chip at a flow rate of 1.1 μl/min. 150 μl ligation mix (1.5X Fast-link ligation buffer (Lucigen, SS000272-D5), 26.7 mM EGTA (bioWorld, 40520008-2), 1.3 mM dNTPs (Thermo Fisher Scientific, R0194), 1.2 mM ATP (Lucigen, SS000391-D3), 16% Glycerol (Sigma-Aldrich, 56-81-5)), 1 ml spacing oil, and 1 ml droplet generation oil were flowed into the device with flow rates of 3 μl/min, 3 μl/min, and 7 μl/min, respectively. Merged droplets were collected into a 1.5 ml tube on ice. The droplets were then exposed to 365 nm UV exposure (160 mJ/cm^2^) for 8 min and incubated at 16 °C for overnight (12-18 h). The droplets were further incubated at 70 °C for 15 min for the inactivation of ligase and end-it enzyme and then broken by adding 100 μl of 100% perfluorooctanol (Sigma-Aldrich, 370533-25G). 15 μl of 20 mM concentration of Proteinase K (Sigma-Aldrich, P2308-100mg) was added to the extracted aqueous phase (∼150 μl) and incubated at 65 °C for 2 h. DNA was first purified using 1.4X SPRIselect bead (Beckman Coulter Life Sciences, B23317) and selected using 1X SPRIselect bead. Finally, DNA was eluted into 20 μl low-EDTA TE buffer (10 mM Tris, 0.1mM EDTA).

### Bisulfite conversion in droplets

Barcoded DNA in 20 μl low-EDTA TE buffer was heated at 95 °C for 1.5 min and immediately put on ice for 2 min to produce single-stranded DNA. The DNA solution was then mixed with 130 μl bisulfite conversion reagent (Promega, N1301, MethylEdge® Bisulfite Conversion System) before being loaded into the bisulfite droplet device (Supplementary Fig. 2) at a flow rate of 5 μl/min, with the oil’s flow rate at 15 μl/min. The collected bisulfite droplets were incubated at 95 °C for 5 min and 54 °C for 1h for bisulfite conversion in droplets. Droplets were then broken to separate the aqueous phase into a new tube. DNA desulphonation and cleanup steps were immediately performed following the manufacturer instructions of the MethylEdge® Bisulfite Conversion System kit from Promega. After the column cleanup, DNA was selected by 1.1X SPRIselect bead and eluted into 11 μl low-EDTA TE buffer. We examined the bisulfite conversion rate using unmethylated lambda phage DNA (Promega, D1521).

### Library preparation

#### Random priming

Treated DNA (11 μl) was mixed with 1 μl of 50 μM random primer (ordered from IDT with the sequence shown in Supplementary Fig. 1) and heated at 95 °C for 3 min and immediately put on ice for 2 min to produce single-stranded DNA. The solution was subsequently mixed with 2 μl blue buffer (10x, Enzymatics, P7010-HC-L), 1 μl of 50 U/μl Klenow exo- (Enzymatics, P7010-HC-L), and 5 μl of 10mM dNTPs on ice and incubated at 4 °C for 5 min, then a slow ramp of 0.1 °C/s to 25 °C, 25 °C for 5 min, then a slow ramp of 0.1 °C/s to 37 °C, and 37 °C for 60 min. DNA was then purified using 1.1x SPRIselect bead and then incubated at 95 °C for 1.5 min and immediately put on ice for 2 min to generate single-stranded DNA, followed by another round purification using 1.1x SPRIselect bead. 19.5 μl low-EDTA TE was used to elute DNA from the SPRIselect beads.

#### Pre-amplification

The 19.5 μl eluate was placed in a 0.2 ml PCR tube preloaded with a PCR reaction mixture containing 25 μl of 2x KAPA HiFi HotStart Uracil+ ReadyMix (Roche, KK2801), 1.5 μl of 25 μM P5 transposon primer (ordered from IDT with the sequence shown in Supplementary Fig. 1), 1.5 μl of 25 μM P7 transposon BS primer (ordered from IDT with the sequence shown in Supplementary Fig. 1), and 2.5 μl DMSO (Sigma, D9170-5VL). The PCR tube was put on a thermocycler and incubated with the program of 98 °C for 30 s, (98 °C for 10 s, 58 °C for 30 s, 72 °C for 30 s) for 5 cycles, and a final 72 °C extension for 1 min. After PCR, DNA was purified using 1x SPRIselect beads and eluted into 19.5 μl low-EDTA TE.

#### Indexing PCR amplification

The 19.5 μl eluate was placed in one microwell of a 96-well plate preloaded with a PCR reaction mixture containing 25 μl of 2x KAPA HiFi HotStart Uracil+ ReadyMix (Roche, KK2801), 1.5 μl of 25 μM N5XX primer (ordered from IDT with the sequence shown in Supplementary Fig. 1), 1.5 μl of 25 μM Ad2.XX primer (ordered from IDT with the sequence shown in Supplementary Fig. 1), and 2.5 μl EvaGreen dye (20x, Biotium, 31000). The 96-well plate was put in a Bio-Rad CFX thermocycler and incubated with the program of 98 °C for 30 s, (98 °C for 10 s, 62 °C for 30 s, 72 °C for 30 s) for 3 cycles to achieve > 200 RFU increase, followed by a final 72 °C extension for 1 min. After PCR, DNA was purified using 0.85x SPRIselect bead and eluted into 20 μl low-EDTA TE.

### Library quantification and sequencing

As shown in Supplementary Fig. 6, the size of the library of 2000 cells was measured using a High Sensitivity D1000 ScreenTape (Agilent Technologies, 5067-5584) on a TapeStation (Agilent Technologies, G2962/A). The concentration of the library was quantified using a KAPA Library Quantification Kit (Roche, KK4824). The library mixed with 5% PhiX or other high-complexity libraries was loaded on a S4 lane of NovaSeq 6000 system. The sequencing followed the standard Illumina protocol with 150 bp paired-end reads. The average number of unique reads per cell generally increases with higher assigned number of reads first until it plateaus (Supplementary Fig. 12).

### Sequencing data analysis

#### Raw read demultiplexing based on barcodes

We extracted single-cell Drop-BS data from raw sequencing reads based on the barcode sequence (25^th^ to 39^th^ base of R2) (Supplementary Code). A barcode whitelist including the barcodes associated with the most raw reads was identified using umi_tools (v1.0.1) whitelist. Sequencing reads with a specific barcode from the barcode whitelist were extracted using umi_tools extract. Sequencing R1 and R2 reads were trimmed by Tim Galore! (v0.6.7) with parameters (--clip_R1 15 --clip_R2 40) to remove artificial bases introduced by end repair during ligation reaction. Trimmed reads were sorted into individual files based on the barcodes using idemp. Alignment to the hg19/mm10 reference genome was performed using Bismark (v0.23.0, bowtie2 (v2.3.5.1)) with the default settings for R2 reads and --pbat for R1 reads (R1 and R2 reads were processed separately).

#### Selection of cell-associated barcodes

We first selected 2.5X (X= the number of cells expected based on experimental parameters) barcodes with the most raw reads. We then created a probability density plot to show the distribution of mapping efficiency for all selected barcodes (Supplementary Fig. 7 and 8). Then we used multiple normal distributions to fit the density plot. The normal distribution with the highest mean mapping efficiency was considered as having our cell-associated barcodes. In order to remove the potential overlap with the adjacent distribution, we used the value of “μ-σ” as the cutoff for mapping efficiency. Thus the mapping efficiency cutoff is specific to each specific dataset.

#### Quality control and construction of allc file for each single cell/barcode

R1 and R2 reads for each single barcode were combined into one file using bamtools merge. Picard (v2.14.0) was used to remove duplicate reads for each barcode. We used methylKit (v1.10.0) in R (v 4.1.2) to identify the methylation state for each cytosine in the unique reads. “allc_*.tsv” file was constructed for each singe cell/barcode, as in previous work^32^.

#### Construction of feature matrix and clustering

We used the allc files to calculate the methylation rate (mCG/CG or mCH/CH) over bins (1 Mb for cell line data to match the low sequencing depth, or 100 kb for the mouse/human brain data to facilitate comparison with previous works^6, 8^) across the genome for each single cell. We combined the methylation rate data from all singe cells into a feature matrix. Single-cell data in the matrix were then normalized by dividing by the global mCG/CG or mCH/CH of the specific single cell. We conducted principal component analysis (PCA using the irlba package (v 2.3.5) in R) to reduce the dimensions of the normalized matrix to the top 50 principal components for Louvain clustering (using the igraph package (v1.3.1), Supplementary Code). Finally, we projected the clustering results using uniform manifold approximation and projection (UMAP (v 0.2.8.0)).

#### CG methylation z-score calculation

We merged unique reads of single cells within each cluster. Then we calculated the CG methylation level of each cluster over previously defined neuron type-specific DMRs^6^. We used scale function in R to calculate the methylation z-scores and plotted the data using pheatmap (v1.0.12) in R.

#### Pseudobulk DMR analysis

Single-cell data within the excitatory neuronal clusters (cluster 1, 4, and 5) from Human Brain 1+2 were merged to generate the excitatory neuronal pseudobulk data. Similarly, the inhibitory neuronal pseudobulk data were produced from the inhibitory neuronal cluster (cluster 3). In these cases, we combined all “allc_*.tsv” files from the excitatory or inhibitory neuronal cells into one “allc_*.tsv” file. For each unique CpG in the combined file, we calculated the total number of methylated cytosines and the total number of cytosines. This information was used as the pseudobulk data for downstream DMR calling. DSS package (v2.48.0) in R was used to call CG-DMRs between the excitatory neuronal data and the inhibitory neuronal data. We further used ChIPseeker package (v1.36.0) in R to annotate the DMRs to genes.

## Data availability

The Drop-BS data on cell lines and mouse brain and the processed Drop-BS data on human brain can be accessed via Gene Expression Omnibus (GEO) under accession number GSE 204691. The raw Drop-BS data on human brain were deposited in dbGaP under accession number phs002123.v2.p1.

## Code availability

Codes used in this study are provided in Supplementary Code.

## Acknowledgements

The study was supported by the United States National Institutes of Health grants R21HG010195 (C.L.), R01GM143940 (C.L.), R01CA243249 (C.L.), R01MH084894 (J.G.-M.), and the Basque Government grant IT-1211-19 (J.J.M.). The authors thank staff members of the Basque Institute of Legal Medicine for their cooperation in the study.

## Author contributions

C.L. and Q.Z. conceived the idea and designed the Drop-BS device and protocol. Q.Z. conducted all Drop-BS experiments and data analysis. S.M. assisted with data analysis. Z.L. carried out the mouse work. B.Z., Z.Z., and G.L. helped with the experimental work. J.J.M. and J.G.-M. provided human brain samples and contributed to data interpretation. Q.Z. and C.L. wrote the manuscript. All authors proofread the manuscript and provided feedback.

