## Supplementary Information for "Droplet-based bisulfite sequencing for high-throughput profiling of single-cell DNA methylomes"

### Supplementary Figures

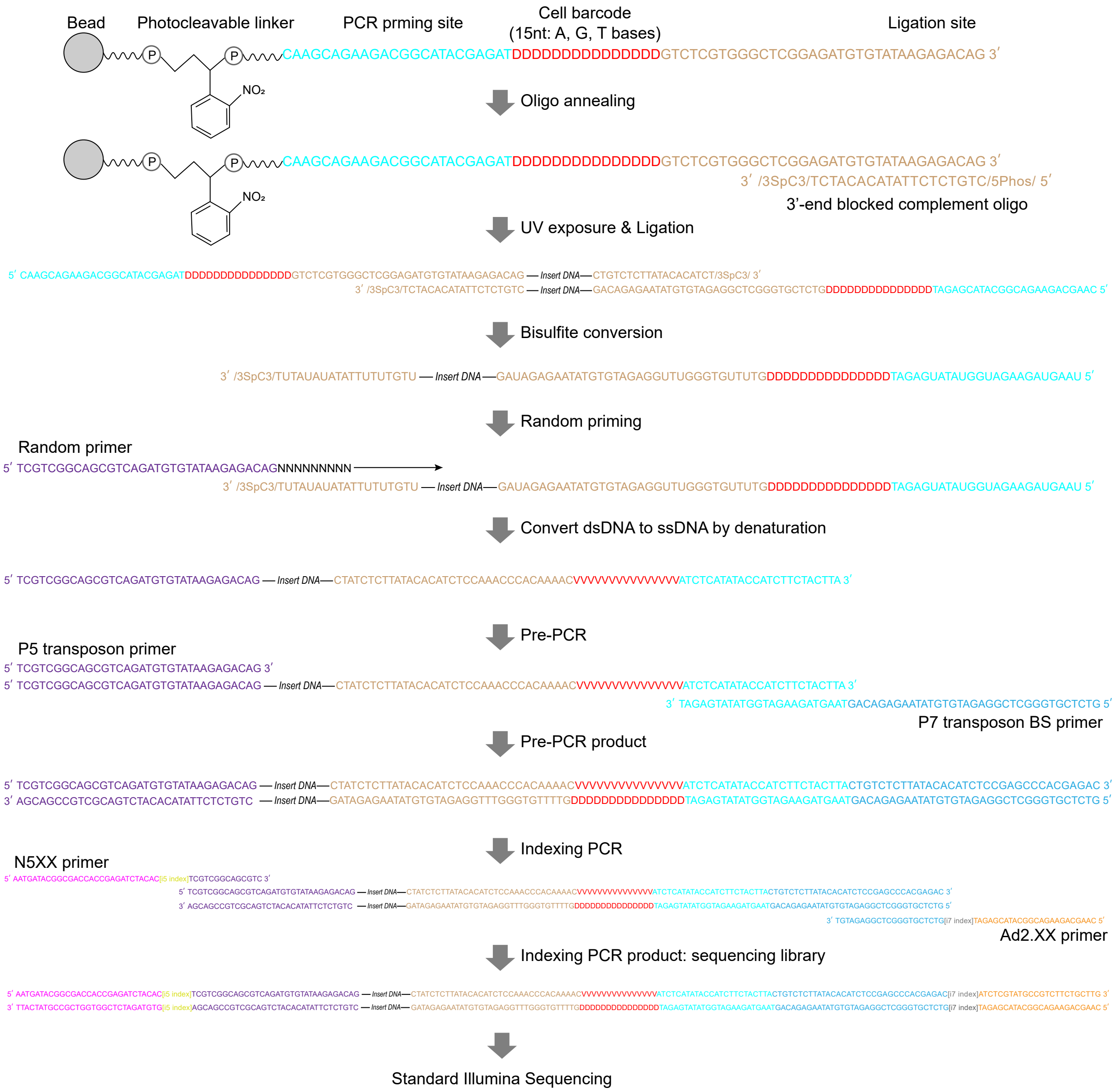


**Supplementary Figure 1** Oligonucleotide and primer sequences involved in the construction of Drop-BS library.

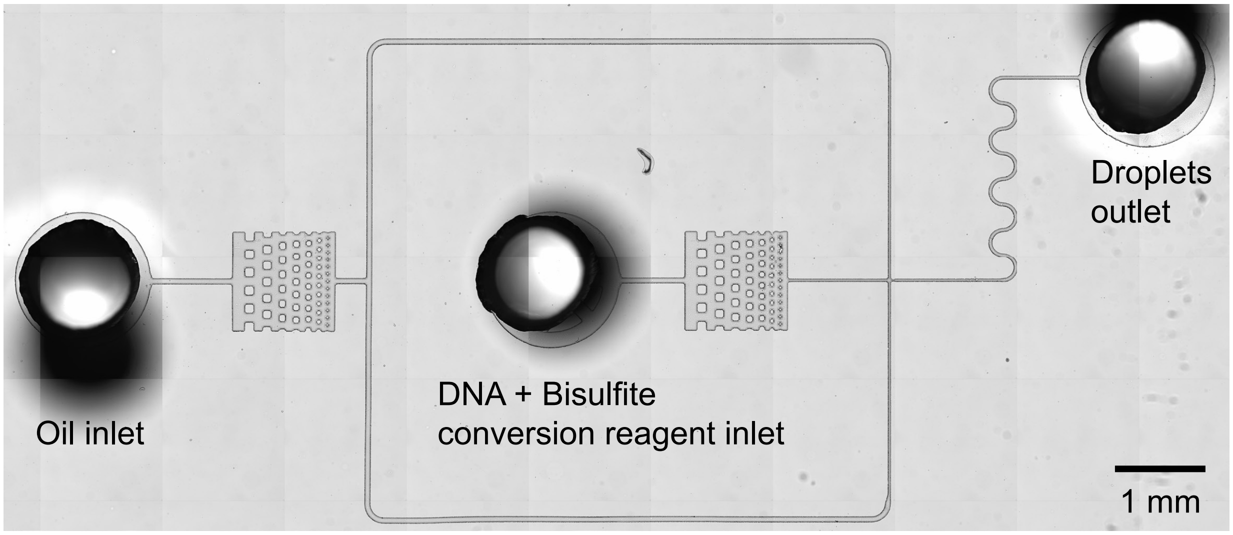


**Supplementary Figure 2** The bisulfite droplet device used in Drop-BS. Filtration structures were placed between the inlets and the narrow channels to trap particles and prevent clogging.

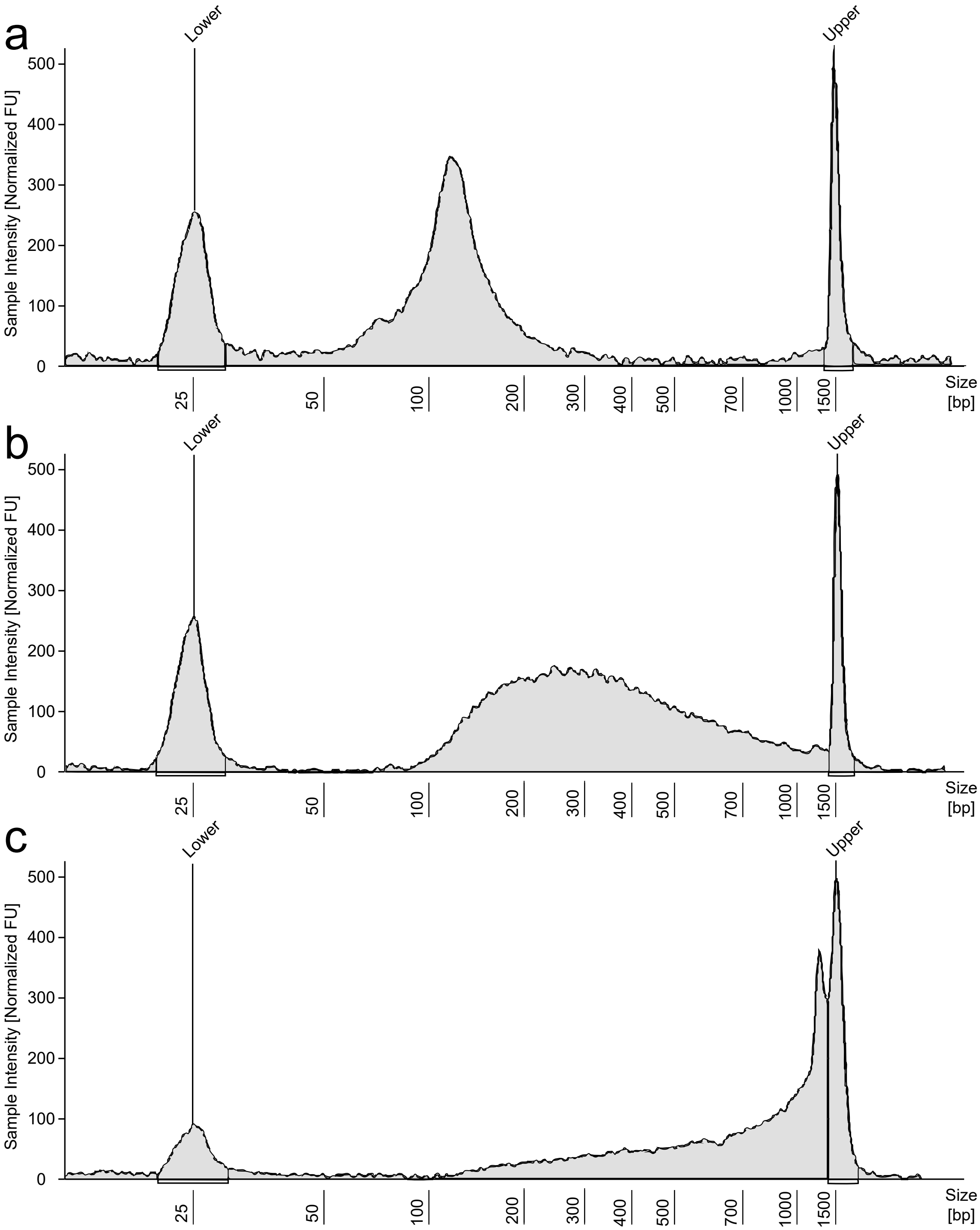


**Supplementary Figure 3** The effect of $CaCl_2$ concentration on the size distribution of single-cell genomic DNA fragmented in droplets. MNase had a concentration of 0.01875 U/µl in droplets in these experiments. (a) 0.25 mM $CaCl_2$; (b) 0.1625 mM $CaCl_2$; (c) 0.0625 mM $CaCl_2$. The final sequencing libraries yielded under these conditions (having a volume of 20 µl) had concentrations of (a) 0.5 nM; (b) 2.5 nM; (c) 0.1 nM, measured by qPCR using the KAPA library quantification kit.

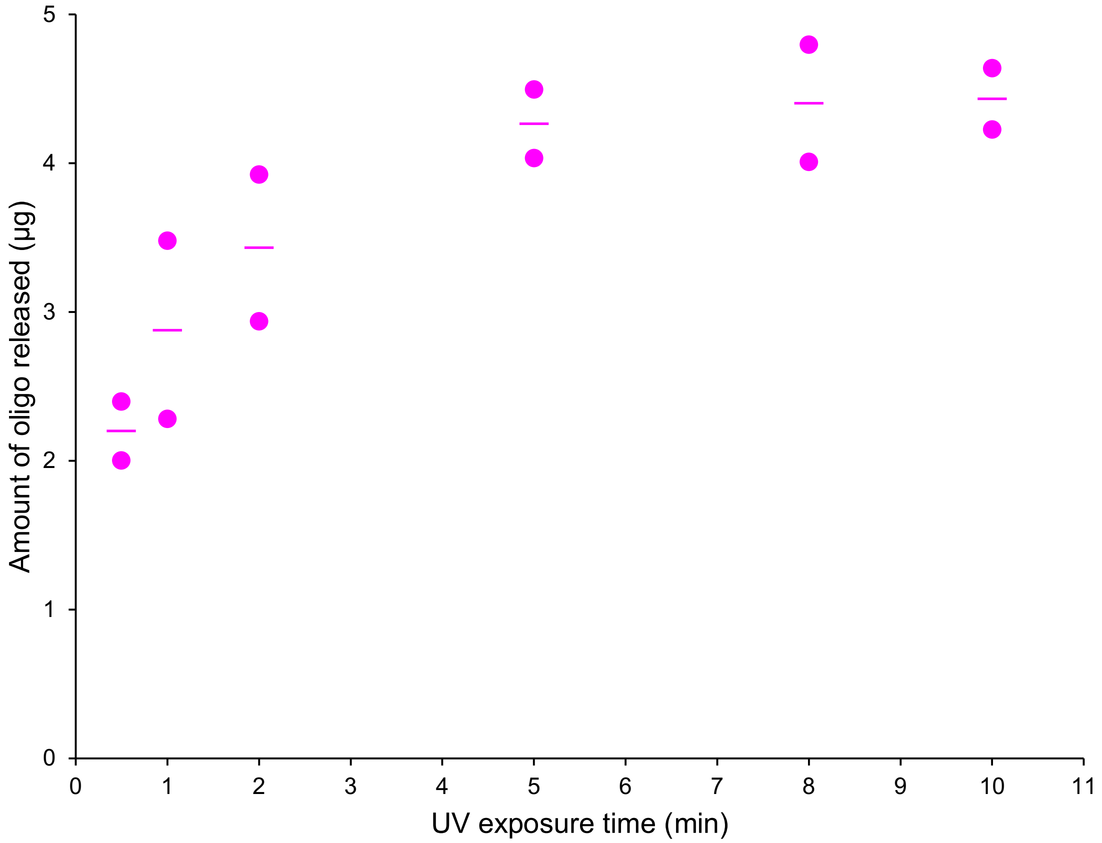


**Supplementary Figure 4** The amount of oligonucleotide released from ~5,000 barcode beads after various UV exposure times. All experiments were conducted in duplicate, and the horizontal lines represent the mean.

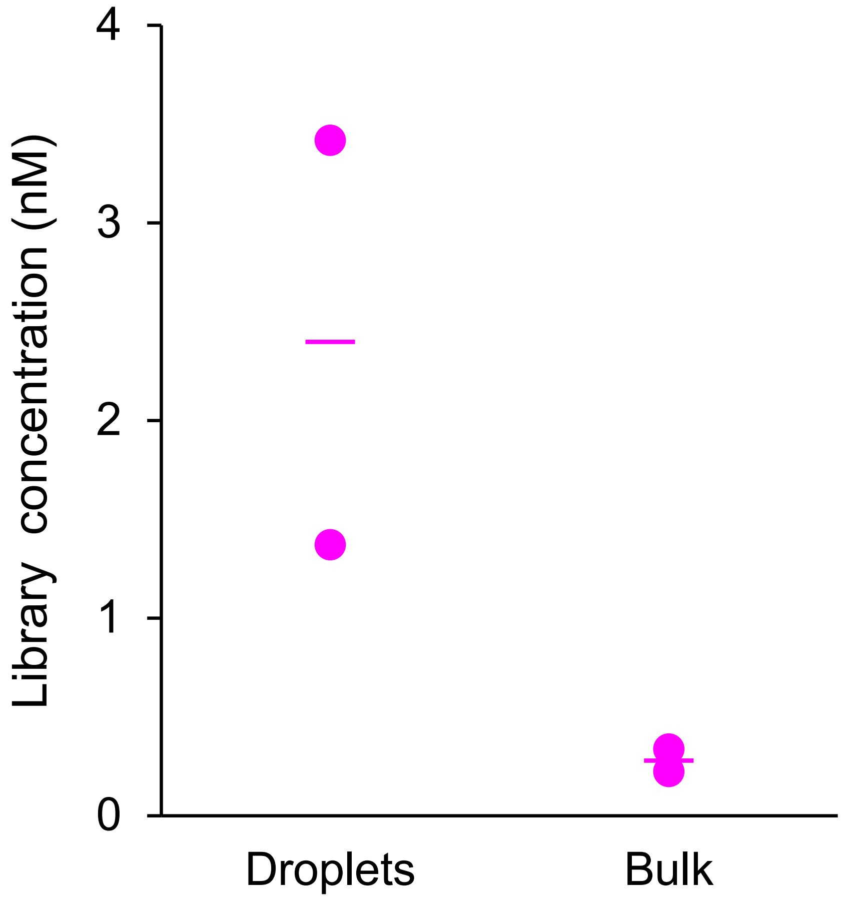


**Supplementary Figure 5** Bisulfite conversion in droplets and bulk (in a tube). We followed Drop-BS protocol to construct these libraries (each starting with ~1,000 GM12878 single cells), except that the “bulk” ones had bisulfite conversion in a tube instead of in droplets. We measured the library concentration using a KAPA Library Quantification Kit (Roche, KK4824). All experiments were conducted in duplicate, and the horizontal lines represent the mean.

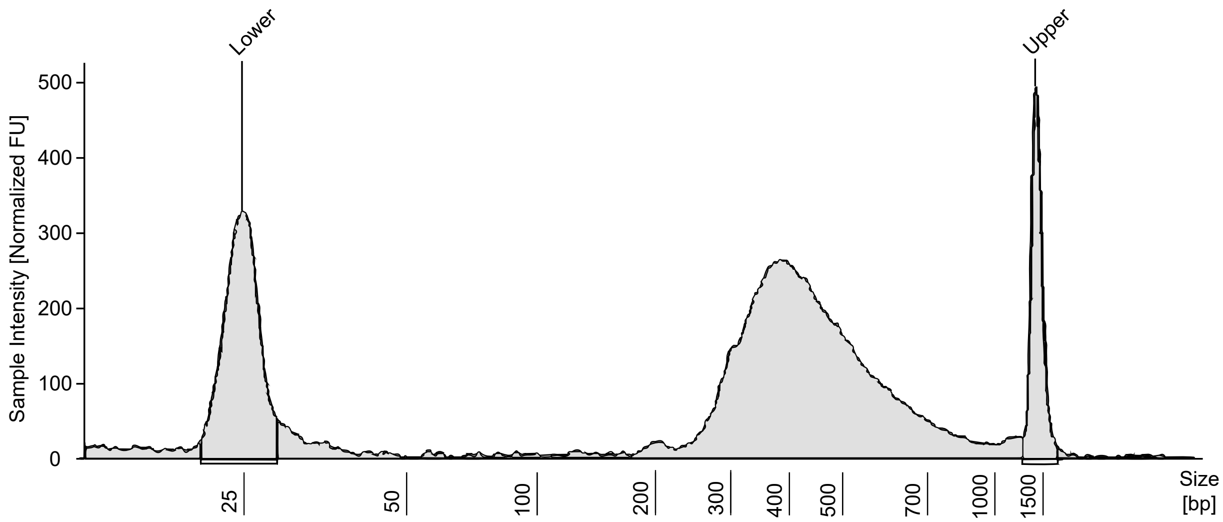


**Supplementary Figure 6** The size profile of a Drop-BS sequencing library measured by an Agilent TapeStation.

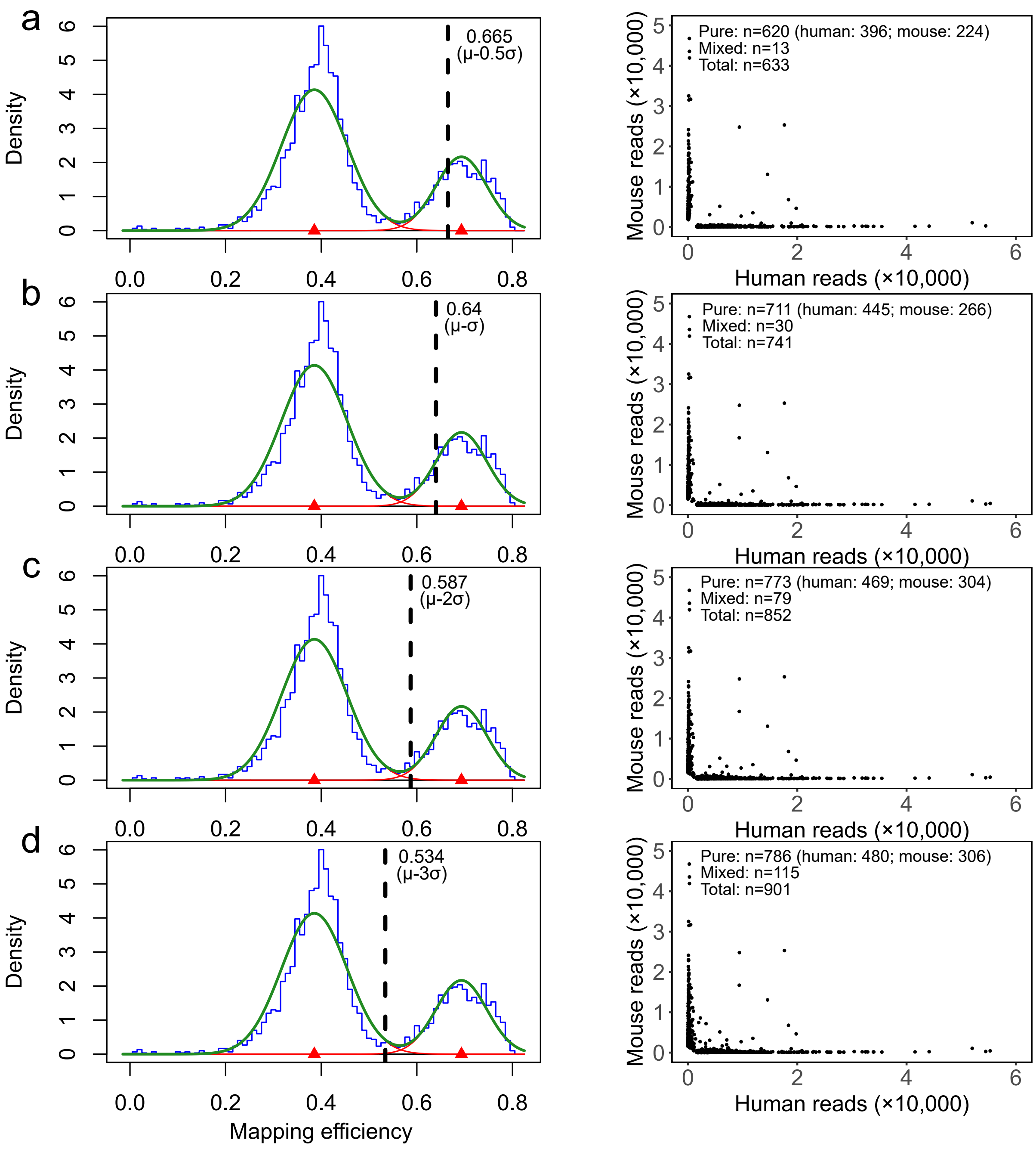


**Supplementary Figure 7** Selection of cell-associated barcodes under various mapping efficiency cutoffs with human/mouse mixed cell sample. GM12878 and mouse brain nuclei were mixed at 1:1 ratio and Drop-BS library of ~1,000 cells were prepared. Each dot is a barcode bearing reads that can align to human genome or mouse genome. "Pure" barcodes refer to the ones with 90% or more of their reads aligned to one genome (human hg19 or mouse mm10). $\mu$ and $\sigma$ are the mean and standard deviation of the fitted normal distribution on the right, respectively. (a) $\mu$ - 0.5$\sigma$; (b) $\mu$ - $\sigma$; (c) $\mu$ - 2$\sigma$; (d) $\mu$ - 3$\sigma$ are used as cutoffs in the probability density plots (left) and the corresponding alignment of selected barcodes to the human and mouse genomes is shown (right). $\mu$ - $\sigma$ is selected as the cutoff that balances the data quality (purity) and the number of cells covered.

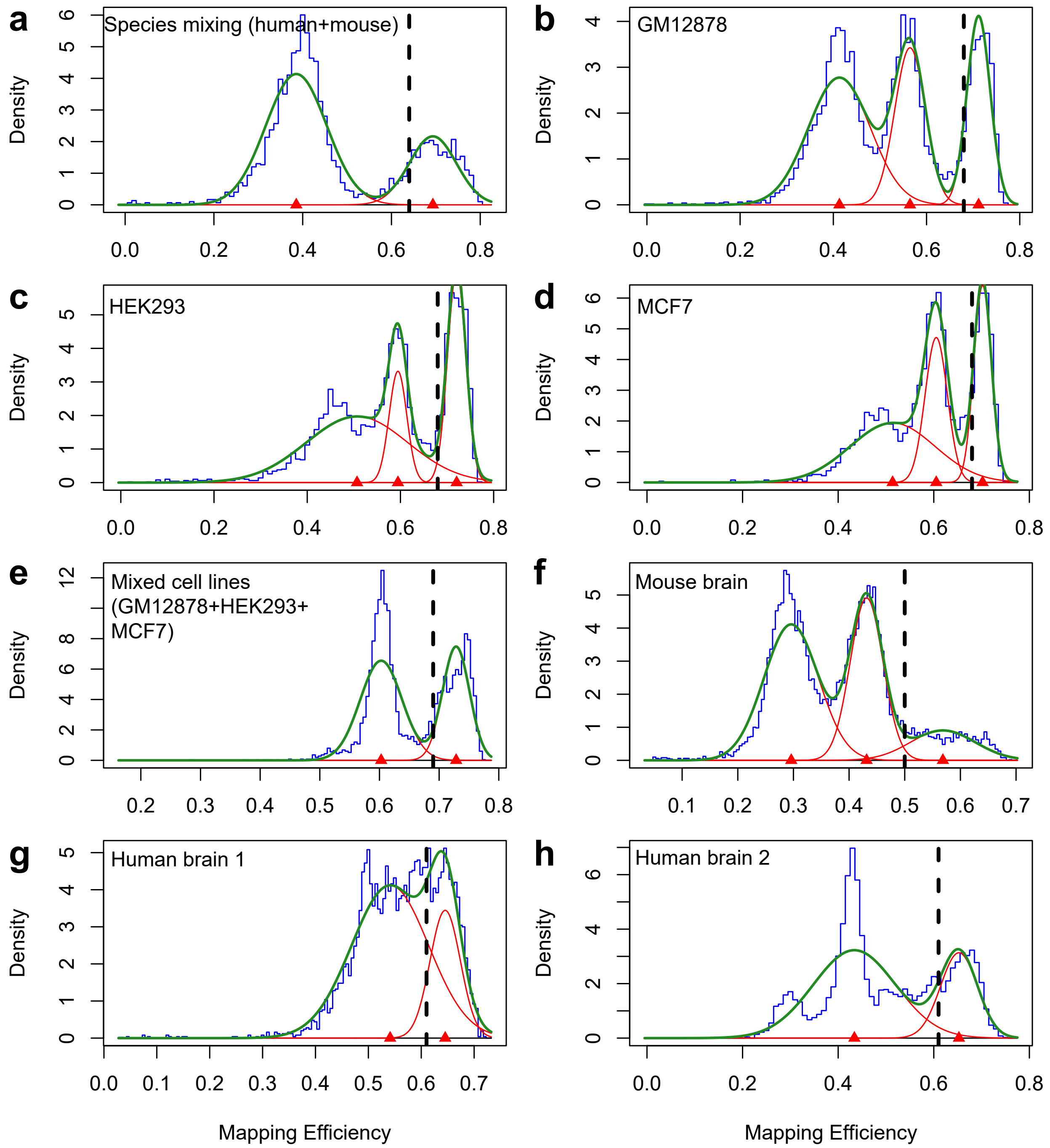


**Supplementary Figure 8** Probability density plots of mapping efficiency for all samples and their corresponding cutoffs ($\mu$-$\sigma$) for selection of cell-associated barcodes. Blue lines: density distributions of mapping efficiency of Drop-BS data. Red lines: fitted normal distributions. Green lines: combined fitted distributions. Black broken lines: the mapping efficiency cutoff, i.e. $\mu$-$\sigma$ of the normal distribution on the right.

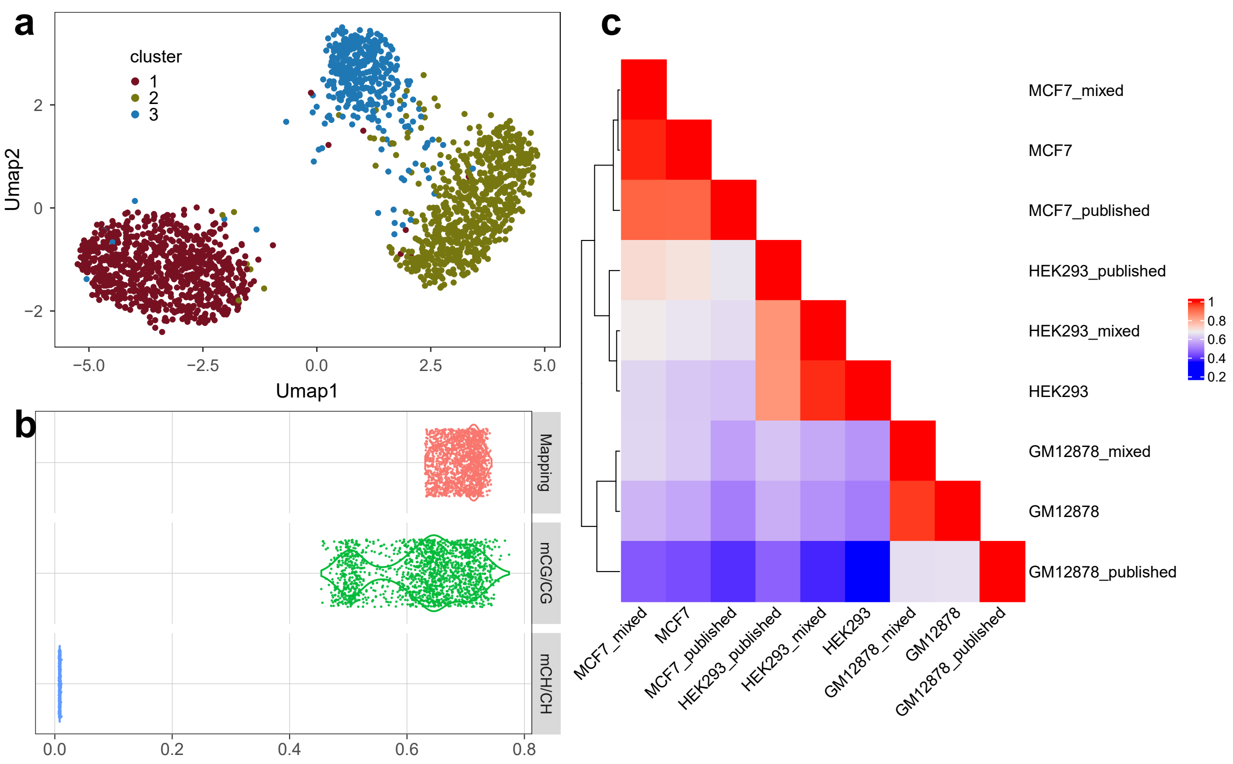


**Supplementary Figure 9** Drop-BS data on cell lines (GM12878, HEK293, and MCF7) and their mixture (containing equal portions of the three cell lines). (a) UMAP visualization of Louvain clustering results on Drop-BS data of a mixture of three human cell lines (Cluster 1: MCF7; 2: HEK293; 3: GM12878). (b) CG methylation rate (mCG/CG), CH methylation rate (mCH/CH), and mapping efficiency of selected high-quality single-cell data from the mixed sample. (c) Pearson correlation and hierarchical clustering among the merged Drop-BS data and published methylomic data based on mCG/CG across equally spaced 1M genomic bins. “GM12878”, “HEK293”, and “MCF7” were the merged single-cell Drop-BS data when each cell line was individually profiled. “GM12878_mixed”, “HEK293_mixed”, “MCF7_mixed” were the merged single-cell Drop-BS data generated on each cell line by clustering the mixed sample. “GM12878_published”, “HEK293_published” and “MCF7_published” were published bulk methylomic data from ENCODE (ENCFF570TIL), GSM1254259, and GSM1328112, respectively.

a

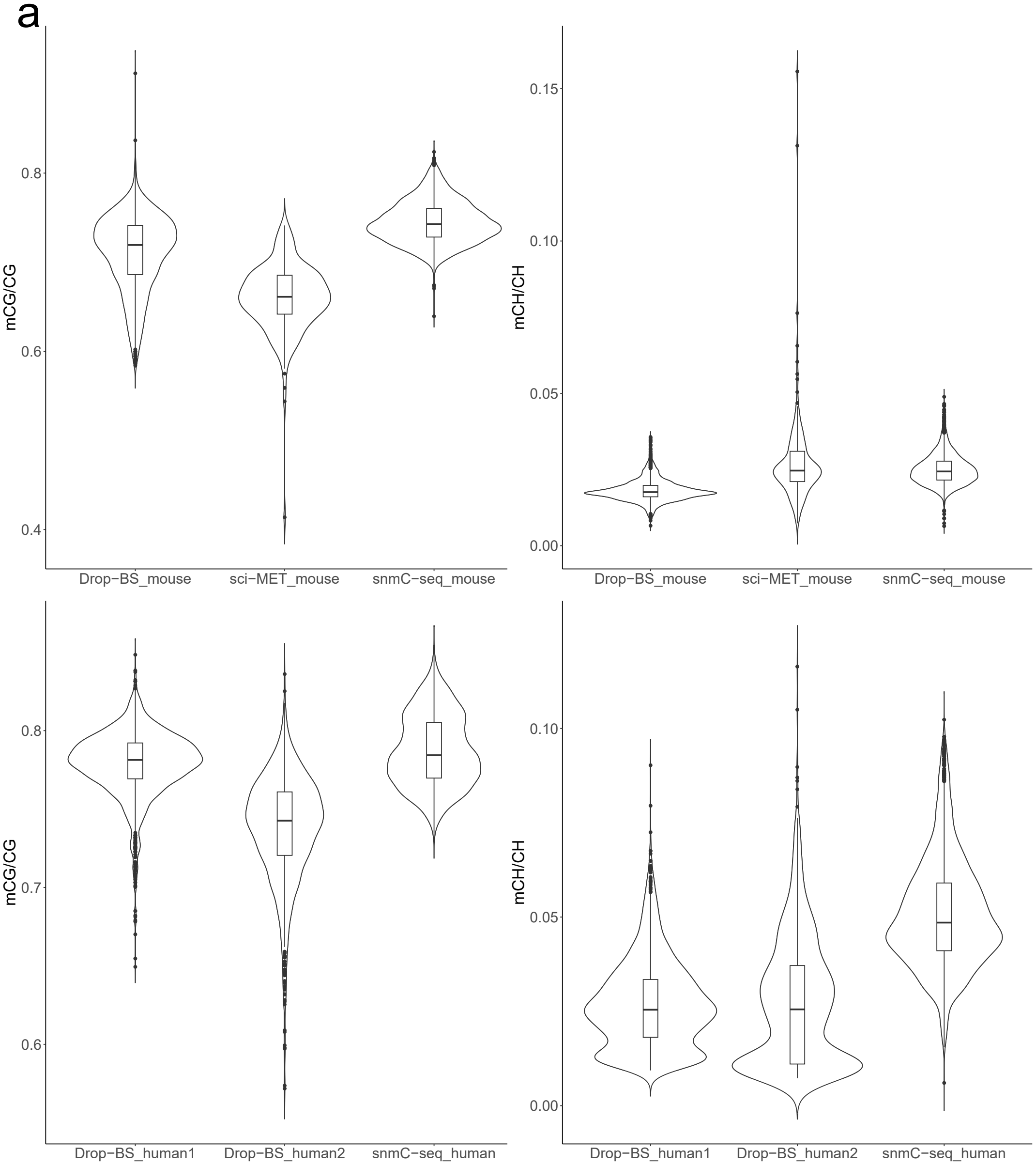


b

| | DropBS mouse brain data | sci-MET mouse brain data | snmC-seq mouse brain data | DropBS human brain 1 data | DropBS human brain 2 data | snmC-seq human brain data |
|---|---|---|---|---|---|---|
| **mCG/CG** | 71.73±3.77% | 65.97±3.82% | 74.49±2.34% | 77.99±2.07% | 73.80±3.39% | 78.71±2.28% |
| **mCH/CH** | 1.85±0.39% | 2.73±1.50% | 2.50±0.49% | 2.70±1.16% | 2.72±1.68% | 5.09±1.44% |

**Supplementary Figure 10** (a) Violin plots and (b) Table of comparison of mCG/CG and mCH/CH of mouse and human brain samples profiled by Drop-BS and other techniques. Drop-BS data are on mouse/human prefrontal cortex tissue. SnmC-seq profiled mouse/human frontal cortex neurons. sci-MET profiled mouse cortex tissue. In the violin plots, the lower and upper hinges correspond to the first and third quartiles. The upper whisker extends from the hinge to the largest value no further than 1.5*IQR from the hinge. The lower whisker extends from the hinge to the smallest value at most 1.5*IQR of the hinge. Data beyond the end of the whiskers are plotted individually.

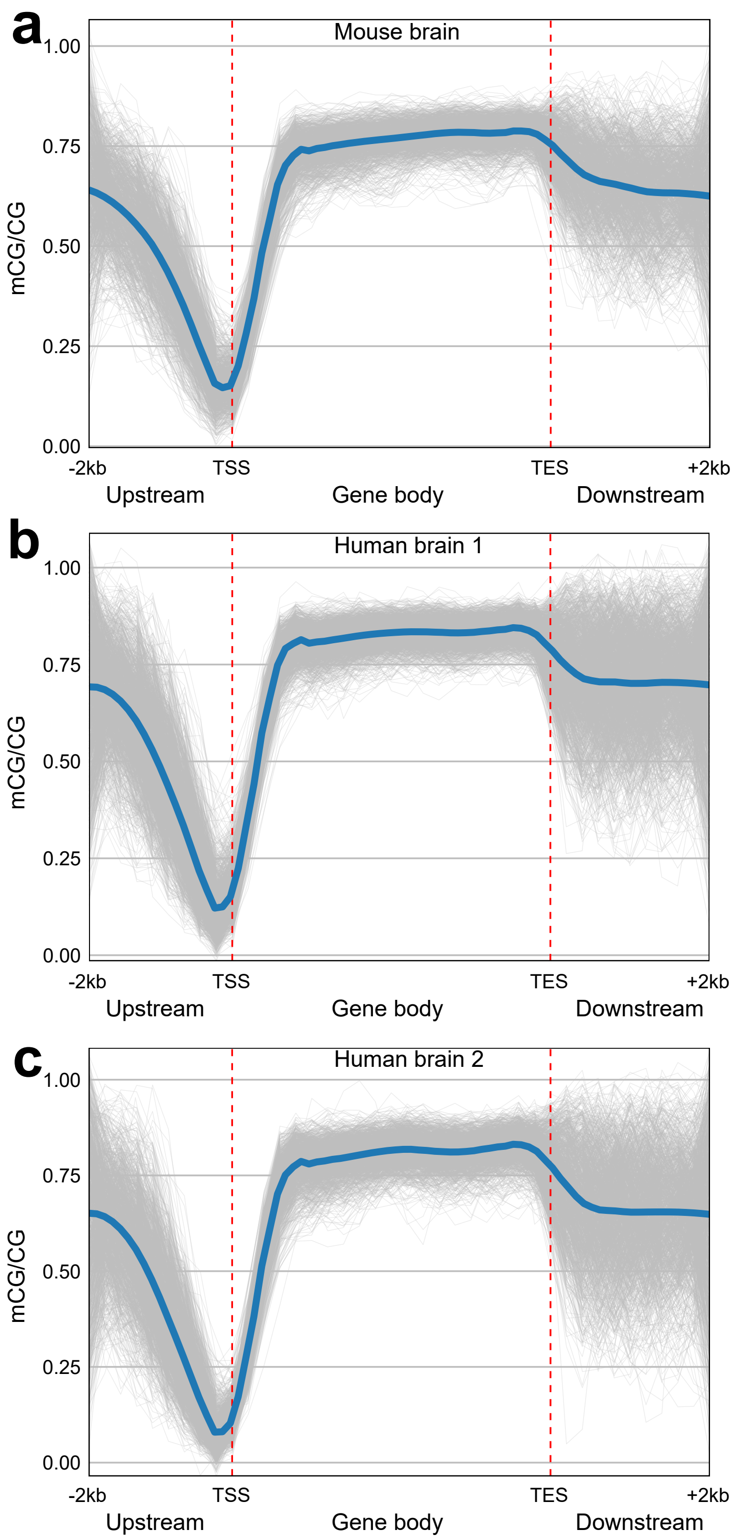


**Supplementary Figure 11** Methylation level (mCG/CG) across upstream 2 kb of TSS (transcription start site), NCBI RefSeqGene body, and downstream 2 kb of TES (transcription termination site) for single cells (grey lines) and an average of all cells (blue line) for mouse PFC sample and human PFC samples.

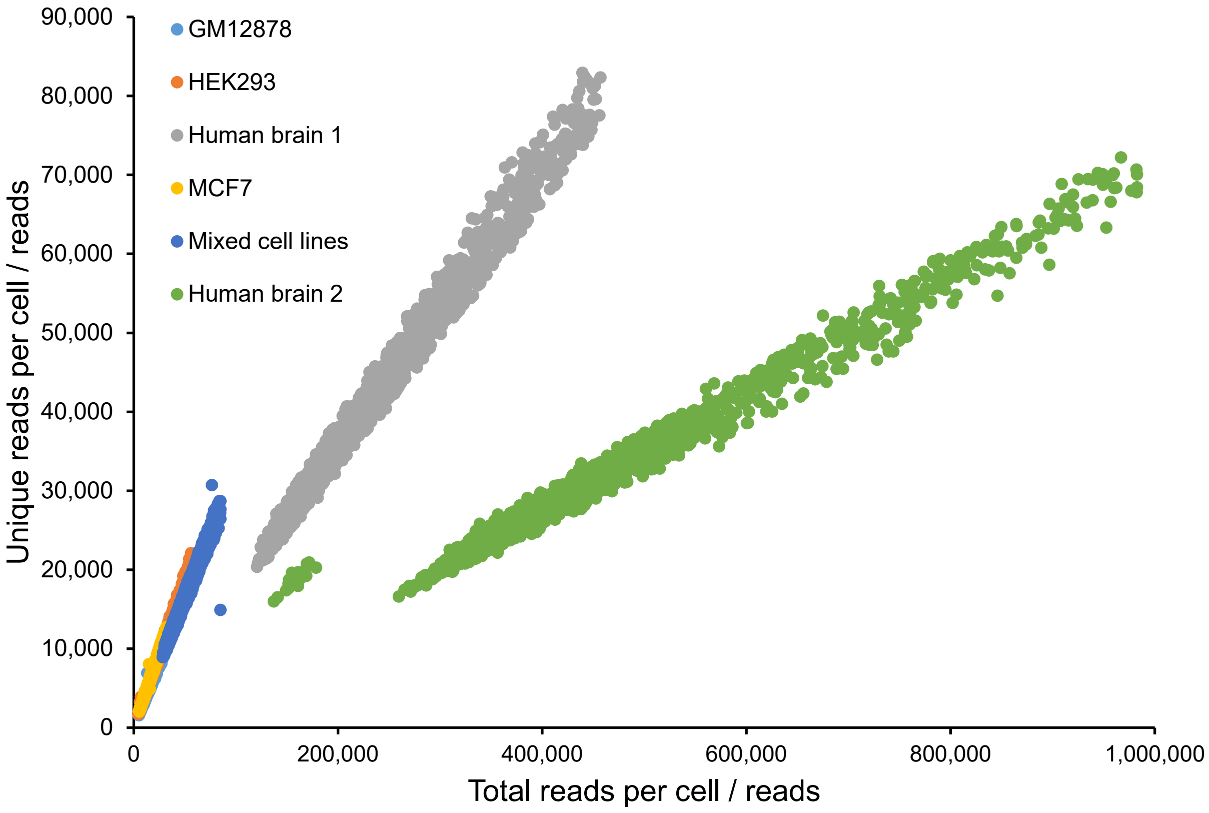


**Supplementary Figure 12** The relationship between the total number of reads and the number of unique reads for each cell in Drop-BS datasets.

**Supplementary Tables**

**Supplementary Table 1.** Metadata for Drop-BS data (excel file attached).

**Supplementary Table 2.** The average mCH/CH of each cluster in combined human brain 1+2 dataset on various functional elements.

| | Cluster 1 | Cluster 2 | Cluster 3 | Cluster 4 | Cluster 5 | Cluster 6 | Cluster 7 |
|---|---|---|---|---|---|---|---|
| Genome wide | 0.0350 | 0.0284 | 0.0126 | 0.0325 | 0.0368 | 0.0266 | 0.0287 |
| CpG Islands | 0.0082 | 0.0081 | 0.0071 | 0.0080 | 0.0078 | 0.0077 | 0.0080 |
| Genic Regions | 0.0318 | 0.0261 | 0.0128 | 0.0296 | 0.0332 | 0.0246 | 0.0265 |
| Intergenic Regions | 0.0378 | 0.0304 | 0.0124 | 0.0350 | 0.0402 | 0.0285 | 0.0306 |
| Promoter Regions | 0.0226 | 0.0197 | 0.0104 | 0.0214 | 0.0228 | 0.0179 | 0.0194 |
| TFBS | 0.0324 | 0.0265 | 0.0124 | 0.0301 | 0.0336 | 0.0247 | 0.0269 |

**Supplementary Table 3.** DMRs and DMR-associated genes identified between excitatory and inhibitory neurons from human brains (excel file attached).

**Supplementary Table 4.** Comparison between the percentages of Drop-BS data (Human brain 1 dataset) on various functional elements and those of the genome occupied by the same elements including CpG Islands (UCSC annotation database), genic regions (NCBI RefSeq), promoter regions (2 kb regions upstream of transcription starting sites of NCBI RefSeq genes), repetitive regions (UCSC annotation database), and TFBS (UCSC annotation database).

| | CpG Islands | Genic Regions | Promoter Regions | Repetitive Regions | TFBS |
|---|---|---|---|---|---|
| Percentage in Drop-BS data | 0.96% | 45.98% | 2.26% | 65.40% | 34.84% |
| Percentage in Genome | 0.39% | 21.73% | 1.04% | 25.29% | 12.83% |

**Supplementary Table 5.** Coverage of genomic regions by each cell in Drop-BS human brain 1 dataset (excel file attached).

**Supplementary Data Sets**

Note: these files can be opened by LayoutEditor that is available free of charge (https://layouteditor.com/).

**Supplementary Data Set 1.** Photomask for the droplet generation device.
**Supplementary Data Set 2.** Photomask for the bisulfite droplet device.
**Supplementary Data Set 3.** Photomask for the droplet fusion device.

**Supplementary Movies**

**Supplementary Movie 1.** Single nuclei encapsulation in the droplet generation device.
**Supplementary Movie 2.** Barcode droplet generation and scDNA droplet reinjection in the droplet fusion device.
**Supplementary Movie 3.** Barcode and scDNA droplets pairing in the droplet fusion device.
**Supplementary Movie 4.** Fusion of scDNA and barcode droplets under dielectrophoresis in the droplet fusion device. The alternating current field was applied via the salt channel electrodes on both sides of the droplet channel.

**Supplementary Code**
Customized codes for data analysis of Drop-BS data.
