## Supplementary Code for "Droplet-based bisulfite sequencing for high-throughput profiling of single-cell DNA methylomes": Code_for_clustering.R

```
 # Code_for_clustering.R is used to perform clustering of single cells.
    # Copyright (C) <2022>  <Sai Ma>

    # This program is free software: you can redistribute it and/or modify
    # it under the terms of the GNU General Public License as published by
    # the Free Software Foundation, either version 3 of the License, or
    # (at your option) any later version.

    # This program is distributed in the hope that it will be useful,
    # but WITHOUT ANY WARRANTY; without even the implied warranty of
    # MERCHANTABILITY or FITNESS FOR A PARTICULAR PURPOSE.  See the
    # GNU General Public License for more details.

    # You should have received a copy of the GNU General Public License
    # along with this program.  If not, see <http://www.gnu.org/licenses/>.

# load package ------------------------------------------------------------
library(umap)
library(irlba)
library(Rtsne)
library(FNN)
library(ggplot2)
umap.defaults$min_dist <- 0.3

options(stringsAsFactors = FALSE)


tsnecols = c("#1F78B4","#68228B","#AAAA44",
             "#60CC52","pink","#DDDD77","#774411","#AA7744","#AA4455","#117744",
             "#000080","#44AA77","#AA4488","#DDAA77")
			 
# load data ---------------------------------------------------------------
rate_bin_ch <- read.table("sample_ID_mC_rate.txt")
#sample_ID_mC_rate.txt: each column is a single cell, each row is a bin, and value is the CH/CG methylation rate.

read_bin_ch <- read.table("sample_ID_mC_read.txt")
#sample_ID_mC_read.txt: each column is a single cell, each row is a bin, and value is the number of CH/CG.

meta <- read.table("sample_ID.tot.txt")
#sample_ID.tot.txt: each row is a single cell. 1st column: the number of mCH; 2nd column: the number of CH; 3rd column: mCH/CH; 4th column: the number of mCG; 5th column: the number of CG; 6th column: mCG/CG;


# filter data -------------------------------------------------------------
# CpG with >10x coverage
filter <- rowSums(read_bin_ch > 10) > 0.3*ncol(read_bin_ch)
print(sum(filter))
rateb_ch <- sweep(rate_bin_ch[filter,],MARGIN=2,FUN="/",STATS=meta[,3]) #for clustering based on mCH/CH
#rateb_ch <- sweep(rate_bin_ch[filter,],MARGIN=2,FUN="/",STATS=meta[,6]) #for clustering based on mCG/CG


# dimension reduction -----------------------------------------------------
run_pca <- function(mat,num_pcs=50,scale=FALSE,center=FALSE){
  set.seed(101)
  SVD = irlba(mat, num_pcs, num_pcs,scale=scale,center=center)
  sk_diag = matrix(0, nrow=num_pcs, ncol=num_pcs)
  diag(sk_diag) = SVD$d
  SVD_vd = t(sk_diag %*% t(SVD$v))
  return(t(SVD_vd))
}

TF_pc <- run_pca(as.matrix(rateb_ch), num_pcs = 50)

umap <- umap(seed=42, t(TF_pc), min_dist=0.1); plot(umap$layout)
tsne <- Rtsne(seed=42, t(TF_pc), pca = F, check_duplicates= T, perplexity = 30, exaggeration=12); plot(tsne$Y) #singles

sumamry <- data.frame(umap1 = umap$layout[, 1],
                      umap2 = umap$layout[, 2],
                      tsne1 = tsne$Y[, 1],
                      tsne2 = tsne$Y[, 2]
                      )
					  
# louvain clustering ------------------------------------------------------
k = 30
knn.pca <- get.knn(t(TF_pc), k = k)
knn.pca = data.frame(from = rep(1:nrow(knn.pca$nn.index), k),
                     to = as.vector(knn.pca$nn.index), 
                     weight = 1/(1 + as.vector(knn.pca$nn.dist)))

suppressWarnings(library(igraph))
nw.norm = graph_from_data_frame(knn.pca, directed = FALSE)
nw.norm = simplify(nw.norm)
lc.norm = cluster_louvain(nw.norm)
unique(membership(lc.norm))
sumamry$cluster <- as.factor(membership(lc.norm))
write.csv(sumamry, "result.csv")

#Plotting
p1 <- ggplot(sumamry, aes(x=umap1, y=umap2, color=cluster)) + geom_point(size=0.8, stroke=0) + theme_bw() + 
  scale_color_manual(values=tsnecols) +
  guides(colour = guide_legend(override.aes = list(size=1)),fill=guide_legend(title="New Legend Title"))+theme_bw() +
  theme(panel.grid.major=element_blank(),panel.grid.minor=element_blank())+
  theme(legend.position = c(0.9,0.3))+
  theme(axis.text = element_text(size = 5))+
  theme(axis.title = element_text(size = 8),
        legend.title=element_text(size=5),
        legend.text=element_text(size=5),
        legend.key.size = unit(0.1, 'cm'))+
  theme(
    panel.background = element_rect(fill='transparent'), #transparent panel bg
    plot.background = element_rect(fill='transparent', color=NA), #transparent plot bg
    panel.grid.major = element_blank(), #remove major gridlines
    panel.grid.minor = element_blank(), #remove minor gridlines
    legend.background = element_rect(fill='transparent'))+ #transparent legend panel
  xlab("Umap1") + ylab("Umap2")+
  theme(axis.title.y = element_text(margin = margin(t = 0, r = 0.2, b = 0, l = 0)))+
  scale_y_continuous(breaks=c(-5,0,5))+
  scale_x_continuous(breaks=c(-2,0,2))

p1
```
