## Supplementary Code for "Droplet-based bisulfite sequencing for high-throughput profiling of single-cell DNA methylomes": Code_to_extract_reads_from_raw_reads.sh

```
#!/bin/bash

#this script is used to generate the barcode whitelist, then extract and write associated reads from raw sequencing reads (named "*R1.fastq.gz" for R1 and "*R2.fastq.gz" for R2) into files ("*R1_extracted.fastq.gz" for R1 and "*R2_extracted.fastq.gz" for R2).
#For more informaiton about umi_tools, refer the umi_tools tutorial on github (https://github.com/CGATOxford/UMI-tools/blob/master/doc/Single_cell_tutorial.md).

F1=`echo *R1*`
R1="${F1%%.*}"

F2=`echo *R2*`
R2="${F2%%.*}"

MD=$PWD

umi_tools whitelist --subset-reads=500000000 --stdin $R2".fastq.gz" --error-correct-threshold=1 --extract-method=regex --bc-pattern='(?P<discard_1>TAAGTAGAAGATGGTATATGAGAT){s<=4}(?P<cell_1>.{15})(?P<umi_1>.{0})' --set-cell-number=5000 --log2stderr > whitelist_1bp_5000_allreads.txt

umi_tools extract --extract-method=regex --bc-pattern='(?P<discard_1>TAAGTAGAAGATGGTATATGAGAT){s<=4}(?P<cell_1>.{15})(?P<umi_1>.{0})' --error-correct-cell 1 --stdin $R2".fastq.gz" --stdout $R2"_extracted.fastq.gz" --read2-in $R1".fastq.gz" --read2-out=$R1"_extracted.fastq.gz" --filter-cell-barcode --whitelist=whitelist_1bp_5000_allreads.txt
```
