## Supplementary Code for "Droplet-based bisulfite sequencing for high-throughput profiling of single-cell DNA methylomes": README.txt

This README is associated with "Code_for_clustering.R".

System requirements:
software dependencies: R (v4.1.2) with irlba package (v2.3.5), igraph package (v1.3.1), umap (v0.2.8.0), Rtsne (v0.16), FNN (v1.1.3.1), ggplot2 (v3.3.6).
operating systems: x86_64-w64-mingw32/x64 (64-bit)

Installation guide:
Software and packages can be installed according to their own descriptions.

Demo:
The dataset for demo is included in this zip file. Instructions to run data analysis are included as comments in "Code_for_clustering.R".
The expected output is Fig. 4a in the manuscript. The expected run time is less than 3 min.

Instructions for use:
Instructions to run the analysis are included as comments in "Code_for_clustering.R".
